# Voltage-dependent activation of Rac1 by Na_v_1.5 channels promotes cell migration

**DOI:** 10.1101/597088

**Authors:** Ming Yang, Andrew D. James, Rakesh Suman, Richard Kasprowicz, Michaela Nelson, Peter J. O’Toole, William J. Brackenbury

## Abstract

Ion channels can regulate the plasma membrane potential (V_m_) and cell migration as a result of altered ion flux. However, the mechanism by which V_m_ regulates motility remains unclear. Here, we show that the Na_v_1.5 sodium channel carries persistent inward Na^+^ current which depolarizes the resting V_m_ at the timescale of minutes. This Na_v_1.5-dependent V_m_ depolarization increases Rac1 colocalization with phosphatidylserine, to which it is anchored at the leading edge of migrating cells, promoting Rac1 activation. A genetically-encoded FRET biosensor of Rac1 activation shows that depolarization-induced Rac1 activation results in acquisition of a motile phenotype. By identifying Na_v_1.5-mediated V_m_ depolarization as a regulator of Rac1 activation, we link ionic and electrical signaling at the plasma membrane to small GTPase-dependent cytoskeletal reorganization and cellular migration. We uncover a novel and unexpected mechanism for Rac1 activation, which fine tunes cell migration in response to ionic and/or electric field changes in the local microenvironment.

## Introduction

Cellular morphological changes and migration play key roles in normal physiological processes, including during embryonic development and tissue repair. On the other hand, dysfunctional migration occurs in many disease processes, including cancer metastasis (Bravo-Cordero, Hodgson, & Condeelis, 2012). In particular, mesenchymal migration, which is dependent on polarization and formation of lamellipodial protrusions at the leading edge of cells, enables carcinoma cells to escape from primary tumors into surrounding tissues (Friedl, Locker, Sahai, & Segall, 2012).

Dynamic and adaptable changes in actin polymerization lead to altered stiffness, elongation and branching, which in turn permits acquisition of a mesenchymal phenotype and formation of lamellipodial protrusions (Krause & Gautreau, 2014). These changes are tightly regulated by a multiplicity of signaling mechanisms which are coordinated by the Rho family of small GTPases (Ridley, 2015). One of the best studied Rho GTPases, Rac1, plays a critical role in regulating lamellipodia formation and migration (Wu et al., 2009). Rac1 cycles between active (GTP-bound) and inactive (GDP-bound) forms, catalyzed by guanine nucleotide exchange factors (GEFs) and GTPase-activating proteins (GAPs), respectively (Marei & Malliri, 2017). Rac1 translocates between cytoplasmic and plasma membrane-bound compartments, with activation predominantly occurring at the plasma membrane (Garcia-Mata, Boulter, & Burridge, 2011). Translocation of Rac1 to the plasma membrane and interaction with phospholipids precede interaction with GEFs and nucleotide exchange/activation (Das et al., 2015). Rac1 is anchored to the inner leaflet of the plasma membrane through its prenylated C-terminus and polybasic motif (van Hennik et al., 2003), which permits interaction with the anionic phospholipids PIP2, PIP3 and phosphatidylserine (Finkielstein, Overduin, & Capelluto, 2006; Heo et al., 2006; Remorino et al., 2017). Localized clustering of PIP2, PIP3 and phosphatidylserine within the plasma membrane thus permits precise spatial and temporal distribution of Rac1 activation (Fairn et al., 2011; Kay, Koivusalo, Ma, Wohland, & Grinstein, 2012; Remorino et al., 2017). Rac1 activation gradients within heterogeneous signaling nanodomains in turn regulate cytoskeletal rearrangement, lamellipodia formation and migration through binding to WASP-family verprolin-homologous (WAVE) proteins and activation of the actin-related protein 2/3 (Arp2/3) protein complex (Ehrlich, Hansen, & Nelson, 2002; Machacek et al., 2009; Remorino et al., 2017; Ridley, 2011). However, despite its pivotal role in migration, the cellular mechanisms regulating local Rac1 clustering at the plasma membrane are still incompletely understood.

Another set of proteins that play a key role in regulating cellular migration is ion channels (Schwab, Fabian, Hanley, & Stock, 2012). Changes in ion channel expression and/or activity in cancer cells regulate the plasma membrane potential (V_m_) and intracellular signaling cascades as a result of altered ion flux (Brackenbury, 2016; Prevarskaya, Skryma, & Shuba, 2018). Motile cancer cells possess a more depolarized V_m_ compared to terminally-differentiated non-cancer cells (Yang & Brackenbury, 2013). The V_m_ is functionally instructive in regulating cell cycle progression (Sundelacruz, Levin, & Kaplan, 2009), proliferation (Blackiston, McLaughlin, & Levin, 2009; Zhou et al., 2015), differentiation (Sundelacruz, Levin, & Kaplan, 2008), cytoskeletal reorganization (Chifflet, Correa, Nin, Justet, & Hernandez, 2004; Chifflet, Hernandez, & Grasso, 2005; Chifflet, Hernandez, Grasso, & Cirillo, 2003; Nin, Hernandez, & Chifflet, 2009; Szaszi, Sirokmany, Di Ciano-Oliveira, Rotstein, & Kapus, 2005), tissue morphogenesis, regeneration and tumorigenesis (Beane, Morokuma, Adams, & Levin, 2011; Beane, Morokuma, Lemire, & Levin, 2013; Cervera, Alcaraz, & Mafe, 2016; Chernet, Adams, Lobikin, & Levin, 2016; Chernet & Levin, 2014; Lobikin, Chernet, Lobo, & Levin, 2012; Lobikin et al., 2015). V_m_ depolarization controls mitogenic signaling by promoting redistribution of phosphatidylserine and PIP2 in the inner leaflet of the plasma membrane, enhancing nanoclustering and activation of the small GTPase K-Ras (Zhou et al., 2015). However, the mechanisms by which V_m_ regulates other behaviors, including morphological changes and migration, and the dependency of V_m_ on altered activity of specific ion channels expressed on non-excitable cells, remain unclear.

An important class of ion channels in the context of cellular migration is the voltage-gated Na^+^ channels (VGSCs). VGSCs contain a pore-forming *α* subunit (Na_v_1.1-Na_v_1.9), together with one or more smaller β subunits (β1-β4) (Brackenbury & Isom, 2011; Catterall, Goldin, & Waxman, 2003). The canonical function of VGSCs is to regulate V_m_ depolarization during the rising phase of action potentials in electrically-excitable cells (Hille, 1992). In addition, VGSCs not only regulate neuronal pathfinding and migration (Brackenbury et al., 2010; Brackenbury, Yuan, O’Malley, Parent, & Isom, 2013; Patel & Brackenbury, 2015), but also regulate the migration of non-excitable cells (J. A. Black & Waxman, 2013). For example, the Na_v_1.5 subtype is upregulated in breast cancer cells where it plays a critical role in promoting cellular migration, invasion and metastasis (Besson et al., 2015; Brackenbury, 2012; Brackenbury, Chioni, Diss, & Djamgoz, 2007; Fraser et al., 2005; Martin, Ufodiama, Watt, Bland, & Brackenbury, 2015; Nelson, Yang, Dowle, Thomas, & Brackenbury, 2015; Nelson, Yang, Millican-Slater, & Brackenbury, 2015; Roger, Besson, & Le Guennec, 2003). In non-excitable cancer cells, Na_v_1.5 carries a small persistent inward Na^+^ current, which would be expected to depolarize the V_m_ at steady state (Roger et al., 2003; Yang et al., 2012). Na_v_1.5 activity promotes motility and invasive behavior through several mechanisms, including MAPK activation (Carrie D House et al., 2010; C. D. House et al., 2015), up-regulation of CD44 expression (Nelson, Yang, Millican-Slater, et al., 2015), and potentiation of Na^+^/H^+^ exchanger type 1 (NHE1) activity leading to cysteine cathepsin activation (Brisson et al., 2013; Brisson et al., 2011; Gillet et al., 2009). Na_v_1.5 also promotes acquisition of an elongate mesenchymal-like phenotype, cortactin phosphorylation and cytoskeletal reorganization (Brisson et al., 2013; Nelson, Yang, Millican-Slater, et al., 2015). Taken together, these data point to an important role for both Na_v_1.5 and V_m_ in regulating morphological changes and migration. An important question is, therefore, how migratory behavior is regulated by electrical changes and Na^+^ flux at the plasma membrane.

Here, we investigated the role of Na_v_1.5 in regulating V_m_ in non-excitable breast cancer cells. We report that Na_v_1.5 promotes steady state V_m_ depolarization, which in turn leads to localized activation of Rac1 at the plasma membrane, promoting morphological changes and migration. This work uncovers a novel and unexpected voltage-dependent mechanism for Rac1 activation, which would fine tune cell migration in response to ionic and/or electric field changes in the local microenvironment.

## Materials and Methods

### Chemicals

Iberiotoxin, NS-1619, tetrodotoxin (TTX) and veratridine were purchased from Alomone Labs. Phenytoin was from Sigma. Ionomycin was from Cayman Chemical. EHT1864 was from Santa Cruz Biotechnology. Drugs were reconstituted as stock solutions according to manufacturer guidelines and diluted directly into culture medium and/or recording solutions at the indicated working concentrations.

### Cell culture

MDA-MB-231 human breast cancer cells were cultured at 37°C, 5 % CO_2_ in Dulbecco’s modified eagle medium (DMEM) supplemented with 5% fetal bovine serum (FBS) and 4 mM L-glutamine. MDA-MB-231 cells are a well-established cell line for examining the functional activity of Na_v_1.5 in regulating cell behavior and they demonstrate robust endogenous expression of this channel (Brackenbury et al., 2007; Brisson et al., 2013; Brisson et al., 2011; Fraser et al., 2005; Nelson, Millican-Slater, Forrest, & Brackenbury, 2014; Roger et al., 2003; Yang et al., 2012). Cells were confirmed to be mycoplasma-free by the 4’,6-diamidino-2-phenylindole (DAPI) staining method (Uphoff, Gignac, & Drexler, 1992). Molecular identity was confirmed by short tandem repeat analysis (Masters et al., 2001).

### RNA interference

A GFP-expressing MDA-MB-231 cell line lacking functional Na_v_1.5 expression was produced previously by transduction of recombinant lentivirus for shRNA targeting Na_v_1.5 (MISSION pLKO.1-puro shRNA transduction particles; Sigma) (Nelson, Yang, Millican-Slater, et al., 2015). Na_v_1.5-shRNA cells and cells non-targeting shControl cells were maintained in G418 (4 μl/ml, Sigma), blasticidin (2 μl/ml, AppliChem) and puromycin (0.1 μl/ml, Sigma).

### Whole-cell patch clamp recording

The whole-cell patch clamp technique was used to record membrane current and voltage from cells grown on glass coverslips for 48-72 h. Voltage- and current-clamp recordings were made using a Multiclamp 700B amplifier (Molecular Devices) at room temperature. Membrane currents and V_m_ were digitized using a Digidata 1440A interface (Molecular Devices), low-pass filtered at 10 kHz and analyzed using pCLAMP 10.4 software (Molecular Devices). In voltage-clamp mode, signals were sampled at 50 kHz, and the series resistance was compensated by 40-60 %. Linear components of leak were subtracted using a P/6 protocol (Armstrong & Bezanilla, 1977). The standard extracellular physiological saline solution (PSS) contained the following components (in mM): 144 NaCl, 5.4 KCl, 1 MgCl_2_, 2.5 CaCl_2_, 5 HEPES and 5.6 glucose adjusted to pH 7.2 with NaOH. For the Na^+^-free PSS, NaCl was replaced with 144 mM N-methyl-D-glucamine (NMDG) or choline chloride (ChoCl) adjusted to pH 7.2 with HCl. The standard intracellular (pipette) solution contained (in mM) 5 NaCl, 145 KCl, 2 MgCl_2_, 1 CaCl_2_, 10 HEPES and 11 EGTA adjusted to pH 7.4 with KOH. Free Ca^2+^ concentration in the solution was calculated using MaxChelator software (Stanford University). Na^+^ current was activated by depolarizing from −120 mV (250 ms) to voltages in the range −80 to +30 mV in 5 mV increments (50 ms).

### Membrane potential recording

The steady-state V_m_ was recorded in I = 0 mode within 5 s of achieving the whole-cell configuration. To monitor the continuous V_m_ in individual cells in response to pharmacological treatments, cells were held in the whole-cell configuration for 6 min, with 60 s in the standard PSS, followed by 150 s in treatment and a further 150 s for washout of treatment. Where relevant, a vehicle treatment control was included in each experiment. The mean V_m_ over the last 5 s in each of the three stages was used to compare the V_m_ data. The V_m_ signals were sampled at 200 Hz. Liquid junction potentials were calculated using the integrated tool in Clampex 10.4. Dose-response data were fitted to a sigmoidal logistic equation

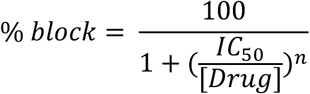

where IC_50_ is the concentration of drug at which half of its maximal effect occurs; and n is the slope giving the Hill coefficient.

### Perforated patch clamp recordings

The perforated patch clamp technique was used to record K_Ca_1.1 currents. The intracellular solution used in perforated patch recording contained (in mM) 5 choline-Cl, 145 KCl, 2 MgCl_2_, 10 HEPES and 1 EGTA adjusted to pH 7.4 with KOH. Nystatin (150 µM) in DMSO was made up and added to the perforated patch intracellular solution on the day of the experiment. Typical series resistance ranged between 20-40 MΩ. K_Ca_1.1 currents were elicited by depolarizing from −120 mV (250 ms) to voltages in the range −60 to +90 mV in 10 mV increments (300 ms). The outward current data were fitted to a single exponential decay (Sanguinetti & Jurkiewicz, 1990)

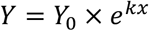

where k is the rate constant.

### Intracellular Na^+^ and Ca^2+^ imaging

Measurement of [Na^+^]_i_ was performed as described in (Roger et al., 2007) with minor modifications. Briefly, 6 × 10^4^ cells grown on glass coverslips for 24 h were labelled with 5 µM SBFI-AM (Sigma) and 0.1 % v/v Pluronic F-127 (Life Technologies) in DMEM with 0 % FBS at 37 °C in the dark for 1 h. Excess SBFI-AM was washed out with 37 °C DMEM supplemented with 5 % FBS. The coverslip was assembled into a RC-20H closed bath imaging chamber (Warner Instruments) and observed at room temperature using a Nikon Eclipse TE200 epifluorescence microscope at 40X. SBFI was alternately excited at 340 and 380 nm, and the fluorescence emission at 510 nm was collected at 8-bit depth using a Rolera-XR Fast 1394 CCD camera (QImaging) controlled by SimplePCI software (Hamamatsu). Calibration of [Na^+^]_i_ was performed after each recording by perfusing two solutions on the cells: 10 and 20 mM Na^+^. They contained (in mM) 149.4 NaCl + KCl, 1 MgCl_2_, 2.5 CaCl_2_, 5 HEPES, 5.6 glucose and 0.02 gramicidin (Sigma), adjusted to pH 7.2 with KOH. In each experimental repeat, [Na^+^]_i_ of ≥ 7 individual cells in the field of view were calculated individually and then averaged. For Ca^2+^ imaging, cells were labelled with 1 µM Fura-2 AM (PromoKine) using the same procedure as above, with an additional wash step using 37 °C phenol red-free DMEM (Life Technologies) after 1 h incubation with the dye, and the images were captured at 20X. In each experimental repeat, the [Ca^2+^]_i_ of ≥ 17 individual cells in the field of view were measured. Each experiment was repeated at least three times.

### Western blotting

Western blotting was performed as described previously (Nelson et al., 2014). The primary antibodies were mouse anti-K_Ca_1.1 (1:500; NeuroMab) and mouse anti-*α*-tubulin (1:10,000; Sigma).

### Cell migration assay

Cell migration was measured using wound healing assays (Yang et al., 2012). Label-free ptychographic microscopy was used to monitor motility in real time (Marrison, Raty, Marriott, & O’Toole, 2013; Suman et al., 2016). Images were acquired over 16 h at 9 min intervals using a Phasefocus VL-21 microscope with an 10X (0.25 NA) lens using 640 nm illumination and equipped with an environmental chamber maintaining the cells at 37 °C in 5% CO_2_. The wound healing experiment was repeated three times on separate days. Image sequences of gap closure were processed using Phasefocus Cell Analysis Toolbox (CAT) software in order to segment and track individual cells at the leading edge and measure wound area. For each image sequence, the following parameters were automatically measured: change in normalized gap area over time; t_1/2_ for gap closure (h), determined by applying a linear fit to the normalized wound area reduction from t = 0 until the time at which the area reduces by 50 %; collective migration (µm/h), defined as:

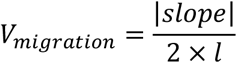

where slope is the rate of area of scratch closure and l is the length of the scratch; instantaneous velocity per cell (µm/s), considering segmented cells with track lengths of ≥ 5 frames; and directionality of leading-edge cells tracked for ≥ 5 frames, relative to the center of the scratch.

### Cell proliferation and invasion assays

Cell proliferation was measured using the thiazolyl blue tetrazolium bromide (MTT) assay, as described (Yang et al., 2012). Cell invasion was quantified using 24-well Corning BioCoat Matrigel Invasion Chambers according to the manufacturer’s guidelines. Briefly, 2.5 × 10^4^ MDA-MB-231 cells were seeded in the upper compartment supplemented with 1 % FBS and appropriate treatments. The lower compartment contained 10 % FBS and appropriate treatments. Cells were incubated at 37 °C for 24 h before the removal of non-invaded cells from the upper chamber using cotton buds. Cells that had invaded through the polyethylene terephthalate membrane were fixed using 4 % paraformaldehyde for 10 min and were washed three times with phosphate buffered saline (PBS), 5 min/time, before staining with DAPI. The membrane was then mounted on a glass microscope slide. DAPI-positive cells on the membrane were viewed at 20X using a Nikon Eclipse TE200 epifluorescence microscope. Experiments were performed on duplicate wells and repeated ≥ 3 times.

### Immunocytochemistry

Cells (1.6 × 10^5^) were cultured on glass coverslips in 4-well plates for 48 h in order to form confluent monolayers. Wounds were made on the coverslip using a P200 pipette tip and debris was washed off with 37 °C DMEM. Appropriate treatments were subsequently applied, and cells were allowed to migrate into the wound for 3 h. This incubation duration was used because it was long enough to show an effect of V_m_ inhibition on migration. Cells were fixed in 4 % paraformaldehyde for 10 min and immunocytochemistry was performed, as described (Nelson et al., 2014). The following primary antibodies were used: rabbit anti-Na_v_1.5 (1:100; Alomone); rabbit anti-K_Ca_1.1 (1:100; Neuromab); mouse anti-Rac1-GTP (1:500; NewEast Biosciences); rabbit anti-total Rac1 (1:10; NewEast Biosciences); mouse anti-CD44 (1:100; AbD Serotec). The secondary antibodies were Alexa 488 or 568-conjugated goat anti-mouse and Alexa 488 or 568-conjugated goat anti-rabbit (1:500; Life Technologies). In some experiments, cells were counter-stained with Alexa 633-conjugated phalloidin (1:50; Life Technologies). In experiments where permeabilized cells were labelled with Alexa 568-conjugated annexin V (1:20; Life Technologies), primary and secondary antibodies were incubated in Ca^2+^-containing annexin V binding buffer (Life Technologies) to preserve annexin V binding. Coverslips were then mounted on glass microscope slides using Prolong Gold with DAPI (Life Technologies). The slides were examined using a Zeiss LSM 710 confocal microscope at 40X. Confocal images were acquired sequentially for each channel. Images (512 x 512 or 1024 x 1024 pixels, dependent on experiment) were initially processed with the Zeiss Zen 2 software, and later exported into ImageJ for analysis. Brightness/contrast was adjusted using the ImageJ “Auto” function.

### Morphology and lamellipodium scoring

Circularity was calculated on manually segmented cells using the free-hand tool in ImageJ (Schneider, Rasband, & Eliceiri, 2012):

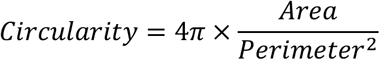

Values approaching 0 indicate a more elongated cell morphology. Feret’s diameter was measured as the maximum distance between any two points along the cell boundary. Three experimental repeats were performed, and 20-25 cells per condition were analyzed for every repeat. To score lamellipodia formation, cells were categorized into two groups: with and without lamellipodium. For each of the three experimental repeats, 40-50 cells per condition were analyzed blinded to treatment.

### Analysis of Rac1 localization in the lamellipodium

The signal densities of F-actin, Rac1-GTP and total Rac1 in the lamellipodium were analyzed in arcs at various distances from the leading edge as described (Dang et al., 2013). The region of interest in the lamellipodia was determined by the dense staining of F-actin. To avoid bias in choosing ROIs, the analysis was carried out blinded to treatment. The ImageJ Radial Profile Extended plugin (Philippe Carl, CNRS, France) was used with starting radius = 0.43 µm, radius increment = 0.43 µm, ending radius = 5.6 µm and total integrated angle = 90°, resulting in 13 arcs in total. Signal densities on each of the arcs were obtained and were normalized to those on the innermost arc. Typically, 20-30 cells at the edge of the wound were analyzed for each condition and the experiment was repeated three times.

### Colocalization analysis

Colocalization of annexin V and Rac1-GTP labeling within regions of interest drawn around the lamellipodium was evaluated using the Coloc2 plugin in ImageJ. The approximate point spread function on acquired images of 1024 x 1024 pixels was 2.71 pixels. For each region of interest, the thresholded Manders M1 and M2 coefficients for annexin V and Rac1-GTP were computed (Manders, Verbeek, & Aten, 1993), together with the Li’s intensity correlation coefficient (Li et al., 2004). Measurements were taken from 30 cells per condition over three experimental repeats.

### In vitro Rac1 activity assay

Cells were cultured to 70 % confluency in 6-well plates prior to addition of pharmacological treatment(s). After incubation for 24 h, total cellular Rac1 activity was evaluated in cell lysates using a colorimetric ELISA-based small GTPase activation assay, according to the manufacturer’s instructions (G-LISA; Cytoskeleton, Inc) (Antonov et al., 2012).

Measurements were obtained from duplicate wells and the experiment was repeated three times.

### Live cell imaging of Rac1 biosensor activation

Cells (1.6 x 10^5^) were cultured in 8-well chamber glass slides. After incubation for 24 h, cells were transfected with 200 ng of a biosensor for Rac1 activation (pTriEx4-Rac1-2G; Addgene plasmid # 66110) (Fritz et al., 2015) using Fugene (Roche) (Nelson et al., 2014). 48 h following transfection, appropriate treatments in phenol red-free DMEM were applied for 3 h prior to imaging. FRET imaging of biosensor activation was performed at room temperature using a Nikon Eclipse TE200 epifluorescence microscope with a 40X (NA 0.60) objective. Acquisition from cells expressing low levels of the biosensor was performed at 8-bit depth using a Rolera-XR Fast 1394 CCD camera (QImaging) controlled by SimplePCI software (Hamamatsu). Acquisition time for donor and FRET channels was typically 200 ms. The donor and acceptor channels were acquired sequentially. The donor, monomeric teal fluorescent protein (mTFP), was excited at 436 nm and emission was collected at 480 nm. FRET emission was collected above 540 nm. The acceptor (Venus) was excited at 480 nm and emission collected above 540 nm. Visualization of cells transfected with a control plasmid expressing mTFP alone, created by site-directed mutagenesis (Nelson et al., 2014), revealed minimal bleed-through in the FRET and Venus channels. Images were taken from ≥49 cells across three experimental repeats. Image processing was performed using ImageJ. Cells were manually segmented using the distribution of biosensor (donor mTFP signal). Images were then background subtracted prior to calculation of the FRET/mTFP integrated density ratio for each cell.

### Statistics

Data are presented as mean ± SEM. GraphPad Prism 7.0d was used to perform all statistical analyses. Polar histograms were plotted using OriginLab Origin 2017. For normally distributed data, paired or unpaired Student’s two-tailed t-test was used to compare two groups, as appropriate. Multiple comparisons were made using 1-way ANOVA or repeated measures ANOVA followed by Tukey *post-hoc* tests, as appropriate. For repeated measures data that were not normally distributed, a Friedman test with Dunn’s *post-hoc* tests was used. Fisher’s exact test was used to examine the distribution of samples within a population. Contingency table data were analyzed using χ^2^ test. Colocalization data were analyzed by 2-way ANOVA. Results were considered significant at P < 0.05.

## Results

### Na_v_1.5 channels depolarize the membrane potential of breast carcinoma cells

The V_m_ of motile cancer cells is generally more depolarized than in terminally-differentiated non-cancer cells (Yang & Brackenbury, 2013). In addition, both depolarized V_m_ and Na_v_1.5 channel activity correlate with metastatic potential (Fraser et al., 2005). However, it is not known whether the relationship between Na_v_1.5, V_m_ and cell behavior is causal. Here, we first investigated whether Na^+^ conductance through plasma membrane Na_v_1.5 channels contributes to the depolarized V_m_ that has been reported previously in MDA-MB-231 cells (Fraser et al., 2005). Na_v_1.5 carries a persistent inward Na^+^ current, small in amplitude compared to the transient Na^+^ current, which plays an important role in shaping the action potential firing pattern, especially in the subthreshold voltage range (George, 2005). In non-excitable cells, this would allow significant accumulation of Na^+^ over an extended period, potentially permitting V_m_ depolarization at steady state. Na_v_1.5 is robustly expressed in MDA-MB-231 cells, both in intracellular compartments and at the plasma membrane (Figure 1A). This agrees with previous reports of VGSC distribution in non-excitable cells (Brisson et al., 2013; Brisson et al., 2011; Carrithers et al., 2009; Carrithers et al., 2007; Fraser et al., 2005; Nelson, Yang, Millican-Slater, et al., 2015; Persson et al., 2014; Yang et al., 2012). We measured Na^+^ currents carried by Na_v_1.5 using whole-cell voltage clamp recording. Depolarization from −120 mV to −10 mV elicited both transient and persistent inward Na^+^ currents (Figure 1B, C). Analysis of the voltage-dependent activation and steady-state inactivation of the Na^+^ current revealed a window current between −50 mV and −10 mV (Figure 1D, E). We next investigated whether Na_v_1.5 could regulate the V_m_. Stable suppression of Na_v_1.5 expression using lentiviral shRNA (Nelson, Yang, Millican-Slater, et al., 2015) significantly hyperpolarized the V_m_ compared to control cells, measured using whole-cell current clamp recording, from −12.6 ± 0.9 mV to −16.0 ± 0.8 mV (P < 0.01; n = 16; t test; Figure 1F). Thus, Na_v_1.5 is required to depolarize the steady-state V_m_.

**Figure 1.**
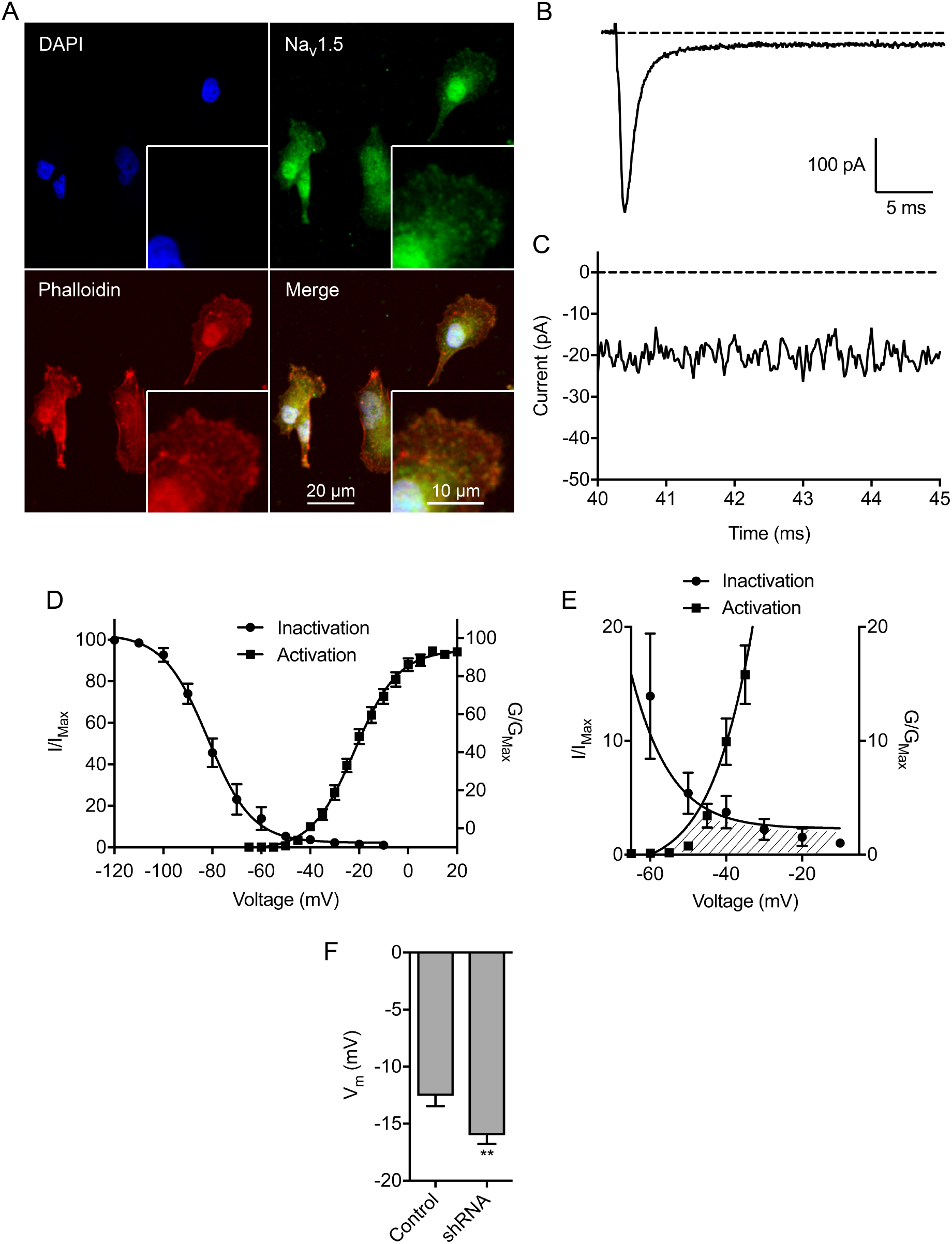
Na_v_1.5 is endogenously expressed in breast carcinoma cells and regulates the membrane potential. (A) MDA-MB-231 cells labeled with Na_v_1.5 antibody (green), phalloidin to label the actin cytoskeleton (red), and DAPI to label the nucleus (blue). Insets show cell peripheries at higher magnification. (B) Typical whole-cell recording showing large transient and small persistent Na^+^ current. The cell was depolarized to −10 mV for 50 ms following a 250 ms pre-pulse at −120 mV. (C) Expanded view of persistent Na^+^ current 40-45 ms following onset of depolarization. (D) Activation and steady-state inactivation of Na^+^ current. Normalized conductance (G/G_Max_) was calculated from the current data and plotted as a function of voltage (n = 14). Normalized current (I/I_Max_) was plotted as a function of the pre-pulse voltage (n = 9). Data are fitted with Boltzmann functions. (E) Expanded view of shaded area under activation and inactivation traces highlighting the window current. (F) Effect of Na_v_1.5 shRNA knock-down on the V_m_ compared to non-targeting shRNA control (n = 16). Data are mean and SEM. *P < 0.05; **P < 0.01; Student’s t-test.

We next evaluated the effect of pharmacologically perturbing Na_v_1.5 activity. The specific VGSC blocker TTX (30 µM) reversibly inhibited the transient and persistent Na^+^ current (P < and P < 0.05, respectively; n ≥ 5; repeated measures ANOVA with Tukey test; Figure 2A-D), consistent with previous reports (Brackenbury et al., 2007; Fraser et al., 2005; Roger et al., 2003). TTX also significantly hyperpolarized the V_m_ from −13.2 ± 1.3 mV to −17.5 ± 1.6 mV (P < 0.01; n = 17; repeated measures ANOVA with Tukey test; Figure 2E, F). On the other hand, the carrier for TTX (148 µM sodium citrate, pH = 4.8) had no effect on the V_m_ (Figure S1A, B). In agreement with the TTX data, the VGSC-inhibiting antiepileptic drug, phenytoin (100 µM) also significantly reduced the transient and persistent Na^+^ current and hyperpolarized the V_m_ (Figure S1C-K). In contrast, veratridine (100 µM), which increases channel open probability by inhibiting inactivation (Ulbricht, 1998), increased the persistent Na^+^ current (Figure 2G, H and Figure S2A, B) and depolarized the V_m_ from −16.37 ± 1.4 mV to −12.6 ± 1.3 mV (P < 0.05; n = 13; t test; Figure 2H). In summary, these data show that pharmacological or genetic inhibition of Na_v_1.5 hyperpolarizes the steady-state V_m_, whereas increasing the persistent Na^+^ current depolarizes the V_m_. Thus, persistent current carried by Na_v_1.5 contributes to V_m_ depolarization.

**Figure 2.**
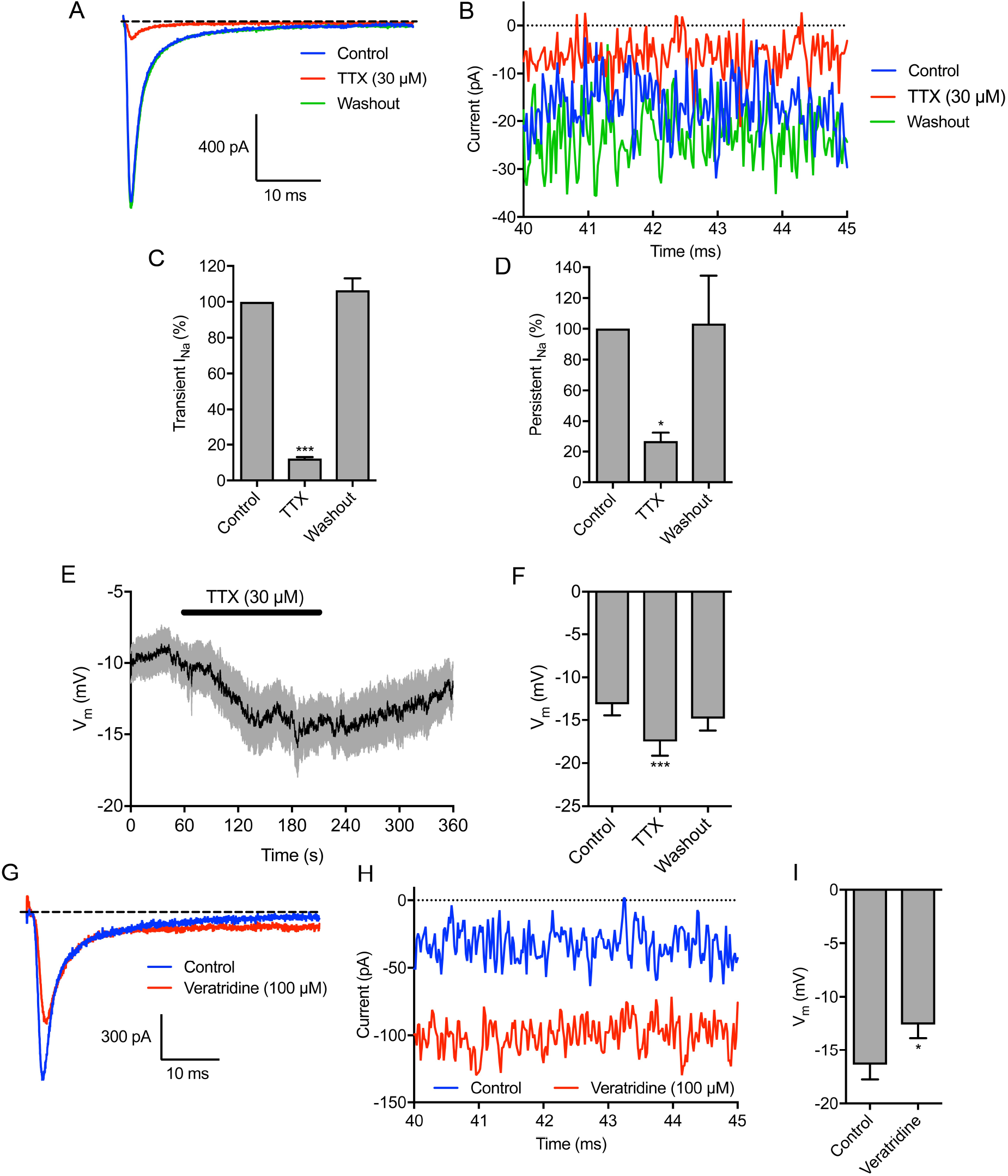
Tetrodotoxin hyperpolarizes, and veratridine depolarizes, the membrane potential. Representative trace showing the inhibitory effect of tetrodotoxin (TTX; 30 µM) on Na^+^ current, and recovery after washout. The cell was held at −120 mV for 250 ms before depolarizing to −10 mV for 50 ms. (B) Expanded view of persistent Na^+^ current 40-45 ms following onset of depolarization. (C) Quantification of the normalized transient Na^+^ current after TTX (30 µM) treatment and following washout (n = 7). (D) Quantification of the normalized persistent Na^+^ current after TTX (30 µM) treatment and following washout (n = 4). (E) V_m_ in control physiological saline solution, after TTX (30 µM) treatment and following washout. Solid line, mean; gray shading, SEM (n = 17). (F) Quantification of V_m_ over the last 5 s in control, TTX, and washout (n = 17). (G) Representative trace showing effect of veratridine (100 µM) on Na^+^ current. The cell was held at −120 mV for 250 ms before depolarizing to 0 mV for 50 ms. (H) Expanded view of persistent Na^+^ current 40-45 ms following onset of depolarization. (I) V_m_ in control PSS and 120 s after perfusion with veratridine (n = 13). Data are mean and SEM. **P < 0.01; ***P < 0.001; repeated measures ANOVA with Tukey test for (C), (D), (F); Student’s t test for (I).

### Membrane potential depolarization is dependent on extracellular Na^+^

To evaluate the sufficiency of extracellular Na^+^ to determine the V_m_, we next removed Na^+^ in the extracellular solution during recording. Replacement of extracellular NaCl with choline chloride reversibly hyperpolarized the V_m_ from −10.4 ± 1.0 to −20.4 ± 2.0 mV (P < 0.001; n = 10; repeated measures ANOVA with Tukey test; Figure 3A, B). Using NMDG to replace extracellular Na^+^ had a similar hyperpolarizing effect (Figure S2C, D). Thus, extracellular Na^+^ is required for steady-state V_m_ depolarization.

**Figure 3.**
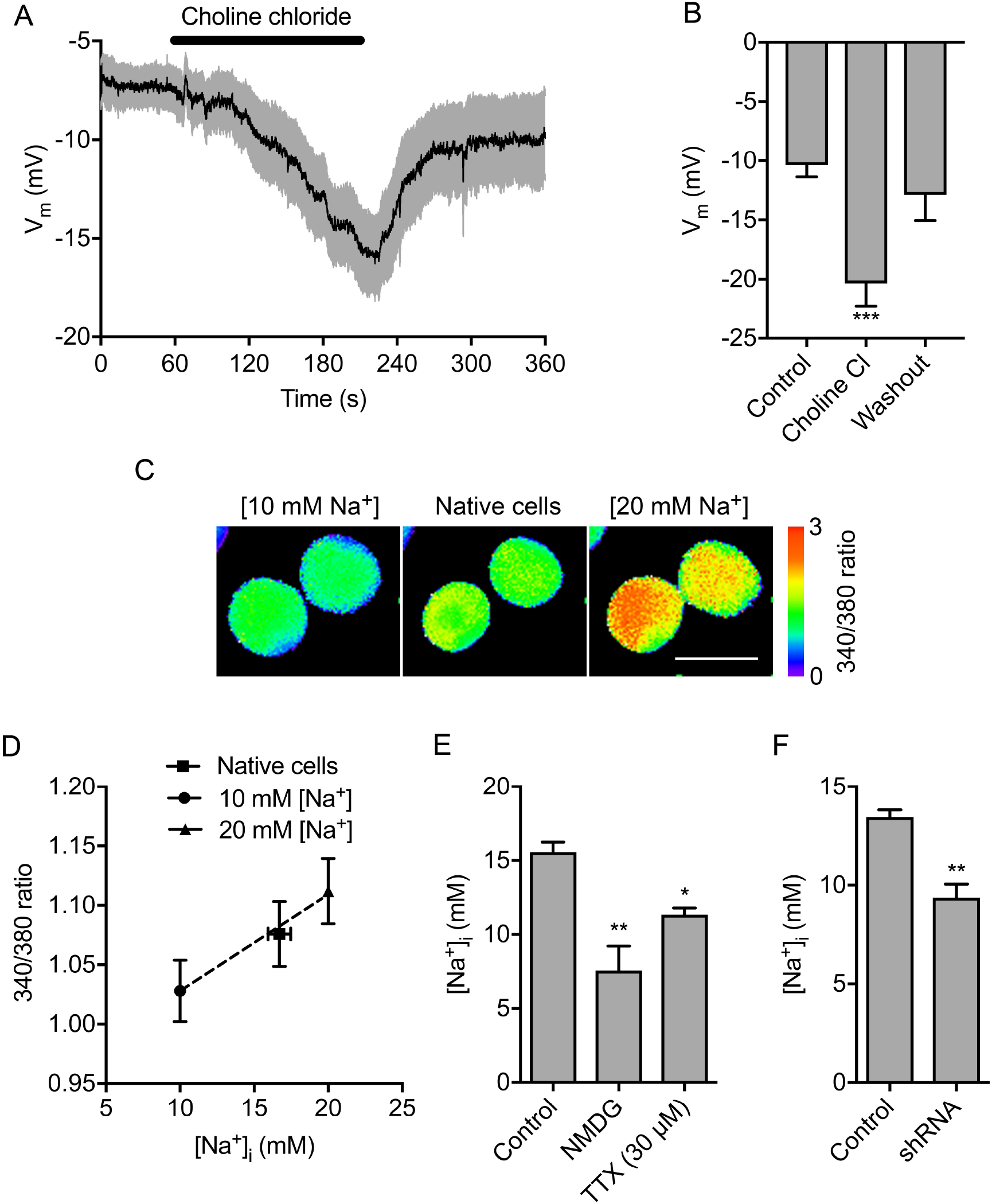
Extracellular Na^+^ sets the membrane potential and intracellular Na^+^ level. (A) V_m_ in control physiological saline solution, after extracellular Na^+^ replacement with choline chloride and following washout. Solid line, mean; gray shading, SEM (n = 10). (B) Quantification of V_m_ over the last 5 s in control, choline chloride, and washout (n = 10). (C) Representative SBFI fluorescence intensity (ratio of emission at 340 nm/380 nm) when cells were perfused with standard physiological saline (center panel), solution containing 10 mM Na^+^ and 20 μM gramicidin (left panel) and solution containing 20 mM Na^+^ and 20 μM gramicidin (right panel). Ratio images are color-coded so that warm and cold colors represent high and low [Na^+^], respectively. Scale bar = 10 μm. (D) Calibration of relationship between SBFI fluorescence intensity (340/380 ratio) and [Na^+^]_i_. Dashed line, linear regression (r^2^ = 0.99; n = 40). (E) Quantification steady-state [Na^+^]_i_ in control physiological saline solution (n = 6), when extracellular Na^+^ was replaced with NMDG (n = 3), and in TTX (30 μM; n = 3). (F) Effect of Na_v_1.5 shRNA knock-down on [Na^+^]_i_ compared to non-targeting shRNA control (n = 3). Data are mean and SEM. *P < 0.05; **P < 0.01; ***P < 0.001; ANOVA with Tukey post hoc test for (A) and (E); Student’s t test for (F).

Inward Na^+^ current through Na_v_1.5 channels would be expected to elevate the intracellular Na^+^ level ([Na^+^]_i_]) at steady state. Indeed, VGSCs have previously been shown to increase [Na^+^]_i_ (Campbell, Main, & Fitzgerald, 2013; Roger et al., 2007). We therefore next investigated the involvement of extracellular Na^+^ in setting [Na^+^]_i_ using the ratiometric Na^+^ indicator SBFI-AM (Figure 3C, D). Replacement of extracellular Na^+^ with NMDG significantly reduced the steady-state [Na^+^]_i_ from 15.6 ± 1.7 mM to 7.6 ± 2.9 mM (P < 0.001; n = 3; ANOVA with Tukey test; Figure 3E). Treatment with TTX also significantly reduced the steady-state [Na^+^]_i_, suggesting that Na_v_1.5 activity contributes to this elevation of [Na^+^]_i_ (P < 0.05; n ≥ 10; ANOVA with Tukey test; Figure 3E). In agreement with this, the [Na^+^]_i_ of Na_v_1.5-shRNA cells was significantly lower than for control cells (9.4 ± 1.2 mM *vs.*13.5 ± 0.6 mM; P < 0.01; n = 3; t test; Figure 3F). On the other hand, Na_v_1.5 inhibition with TTX had no effect on [Ca^2+^]_i_ (Figure S3A-D). Together, these data suggest that Na^+^ influx through Na_v_1.5 channels increases [Na^+^]_i_, but not [Ca^2+^]_i_.

### K_Ca_1.1 regulates the membrane potential but not the Na^+^ level

Na_v_1.5 channels enhance cell migration, invasion, tumor growth and metastasis (Fraser et al., 2005; Nelson, Yang, Millican-Slater, et al., 2015; Roger et al., 2003). Given that Na_v_1.5 depolarizes the steady-state V_m_, and that V_m_ depolarization is functionally instructive in regulating cellular behavior (Yang & Brackenbury, 2013), we reasoned that Na_v_1.5 may regulate cell migration via setting the V_m_. Thus, we sought a means by which to manipulate the V_m_ independent of Na_v_1.5 in order to separate the effects of V_m_ and [Na^+^]_i_ on downstream signaling. To do this, we took advantage of ubiquitously expressed large conductance Ca^2+^-activated K^+^ channel, K_Ca_1.1 (Contreras et al., 2013). We verified that K_Ca_1.1 was robustly expressed in MDA-MB-231 cells (Khaitan et al., 2009; Ma et al., 2012; Roger, Potier, Vandier, Le Guennec, & Besson, 2004), and its expression was unaffected by Na_v_1.5 downregulation (Figure 4A, B). Next, using the perforated patch clamp mode to maintain endogenous [Ca^2+^]_i_, we found that the K_Ca_1.1 opener NS-1619 (1 µM) (Macmillan, Sheridan, Chilvers, & Patmore, 1995) increased outward K^+^ current, confirming K_Ca_1.1 activity (Figure 4C, D). Co-application of the K_Ca_1.1 channel blocker iberiotoxin (100 nM) inhibited the potentiating effect of NS-1619 on the outward current (Figure 4C, D), confirming specificity of NS-1619. We next studied the effect of NS-1619 on the V_m_. NS-1619 hyperpolarized the steady-state V_m_ in a dose-dependent manner (Figure 4E). Importantly, 1 µM NS-1619 hyperpolarized the V_m_ by 4.4 mV from −10.6 ± 1.3 mV to −15.0 ± 0.6 mV (P < 0.01; n ≥ 12; t test; Figure 4F). This shift is equivalent to the hyperpolarization elicited by inhibiting Na_v_1.5 activity (Figure 2F, I). In contrast, NS-1619 treatment did not significantly alter the [Na^+^]_i_ (P = 0.93; n = 22; paired t test; Figure 4G) or [Ca^2+^]_i_ (Figure S3E, F). Finally, the V_m_ recorded using the standard intracellular solution (free [Ca^2+^] = 5.7 nM) was not significantly different from that recorded using intracellular solution with 100 nM free [Ca^2+^] (P = 0.82; n = 12; t-test; Figure 4H), suggesting that at physiological [Ca^2+^]_i_ (Sareen, Darjatmoko, Albert, & Polans, 2007; Winnicka, Bielawski, Bielawska, & Surazynski, 2008), K_Ca_1.1 channels do not contribute to regulating the V_m_ in the absence of NS-1619. In summary, activation of K_Ca_1.1 hyperpolarizes the V_m_ to the same extent as Na_v_1.5 inhibition, thus providing an experimental means by which to manipulate the V_m_ independent of Na^+^.

**Figure 4.**
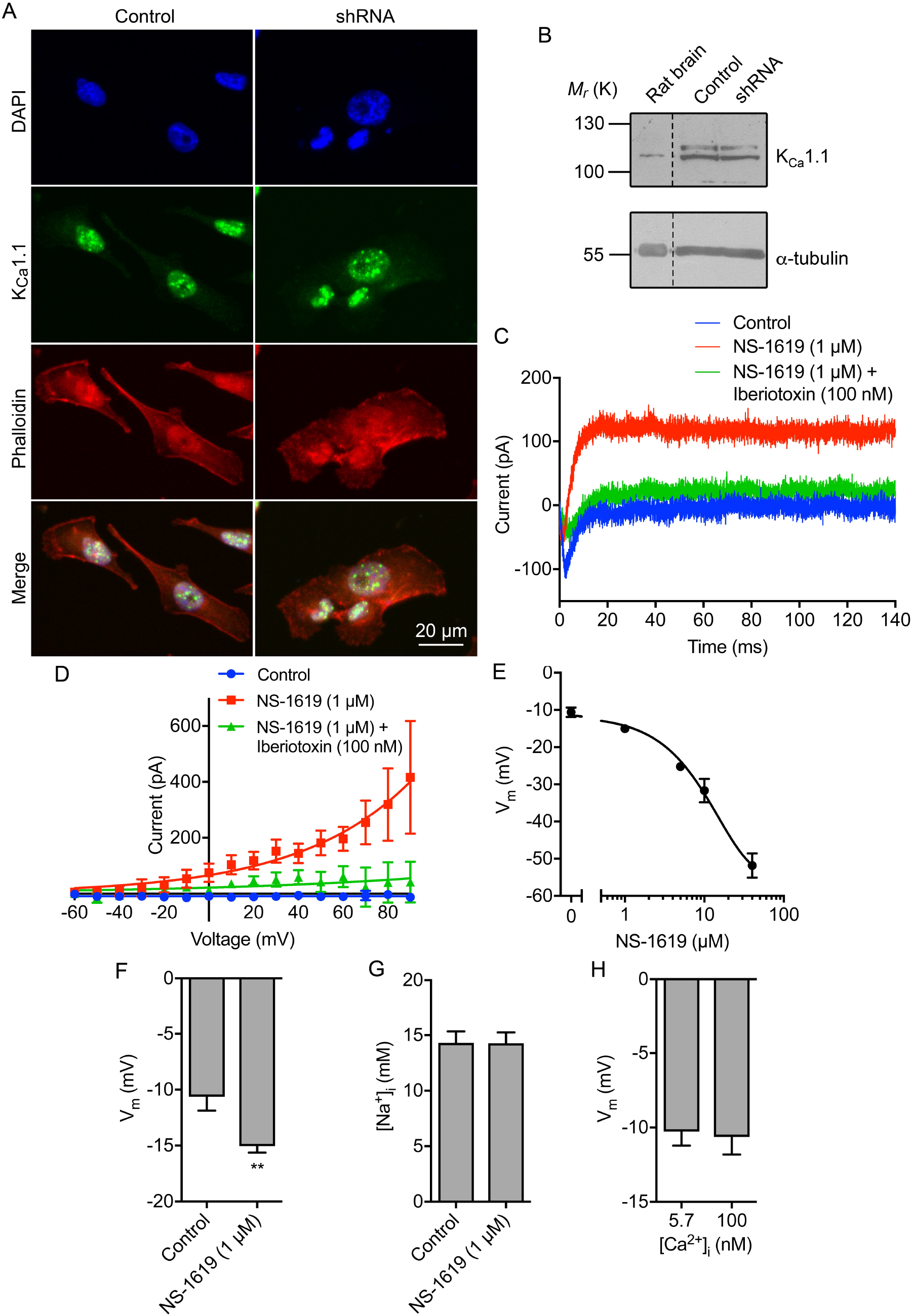
The large conductance Ca^2+^-activated K^+^ channel K_Ca_1.1 regulates the membrane potential but not intracellular Na^+^. (A) MDA-MB-231 cells labeled with K_Ca_1.1 antibody (green), phalloidin to label the actin cytoskeleton (red), and DAPI to label the nucleus (blue). Western blot of K_Ca_1.1 in control MDA-MB-231 cells and cells in which Na_v_1.5 has been knocked down with shRNA. Positive control = rat brain lysate. Loading control = *α*-tubulin. Representative perforated patch clamp recording showing activation of outward current using the K_Ca_1.1 activator (NS-1619; 1 µM) and inhibition with iberiotoxin (100 nM). The cell was held at −120 mV for 250 ms before depolarization to +60 mV for 300 ms. (D) Current-voltage relationship of the K_Ca_1.1 current. Cells were held at −120 mV for 250 ms before depolarization to voltages ranging from −60 to +90 mV in 10 mV steps for 300 ms (n = 5). Data are fitted with single exponential functions. (E) Dose-dependent effect of NS-1619 on the steady-state V_m_ (n ≥ 6). Data are fitted to a sigmoidal logistic function. (F) Effect of NS-1619 (1 µM) on steady-state V_m_ (n = 12). (G) Effect of NS-1619 (1 µM, 5 min) on [Na^+^]_i_ (n = 22). (H) V_m_ recorded using intracellular solution with free [Ca^2+^] buffered to 5.7 nM *vs.* 100 nM (n ≥ 10). Data are mean and SEM. **P < 0.01; Student’s paired t-test.

### Na_v_1.5-dependent membrane potential depolarization regulates cell migration and morphology

Using NS-1619 and TTX as tools to modify the V_m_ independent of, or as a result of Na_v_1.5 activity, respectively, we next investigated the effect of V_m_ depolarization on cell migration in wound healing assays using label-free ptychographic imaging (Figure 5A). Both TTX and NS-1619 slowed the rate of wound closure (Figure 5B). The t_1/2_ of wound closure increased from 5.7 ± 1.1 h to 9.8 ± 0.7 h in the presence of TTX and 9.9 ± 0.1 h in the presence of NS-1619 (P < 0.05; n ≥ 5; ANOVA with Tukey test; Figure 5C). Similarly, the collective migration of cells moving into the wound was reduced from 7.8 ± 0.7 µm/h to 5.5 ± 0.4 µm/h in the presence of TTX and 4.9 ± 0.5 µm/h in the presence of NS-1619 (P < 0.05; n ≥ 5; ANOVA with Tukey test; Figure 5D). The instantaneous velocity of individual cells at the leading edge was also reduced from 6.2 x 10^-3^ ± 5.6 x 10^-5^ µm/s to 4.7 x 10^-3^ ± 5.0 x 10^-5^ µm/s and 4.4 x 10^-3^ ± 4.5 x 10^-5^ µm/s, for TTX- and NS-1619-treated cells, respectively (P < 0.001; n ≥ 2662; Kruskal-Wallis with Dunn’s test; Figure 5E). In addition, both TTX and NS-1619 caused a small, but statistically significant, disruption in the directionality of cells at the leading edge, reducing the proportion of cells migrating directly into the wound (P < 0.001; Friedman with Dunn’s test; Figure 5F). Hyperpolarizing the V_m_ with NS-1619 did not significantly alter cell proliferation measured in an MTT assay (Figure S4A). Given that inhibiting Na_v_1.5 with TTX has no effect on the proliferation of MDA-MB-231 cells (Fraser et al., 2005; Gillet et al., 2009; Roger et al., 2003), together, these data suggest that both Na_v_1.5 activity and V_m_ depolarization promote cellular migration, without affecting proliferation. On the other hand, whilst TTX significantly reduced invasion into Matrigel, consistent with previous reports (Brackenbury et al., 2007; Driffort et al., 2014; Fraser et al., 2005; Nelson, Yang, Millican-Slater, et al., 2015; Roger et al., 2003; Yang et al., 2012), NS-1619 had no effect (Figure S4B, C). Thus, Na_v_1.5 promotes cellular migration via V_m_ depolarization, whereas Na_v_1.5-dependent promotion of invasion occurs via a V_m_-independent mechanism.

**Figure 5.**
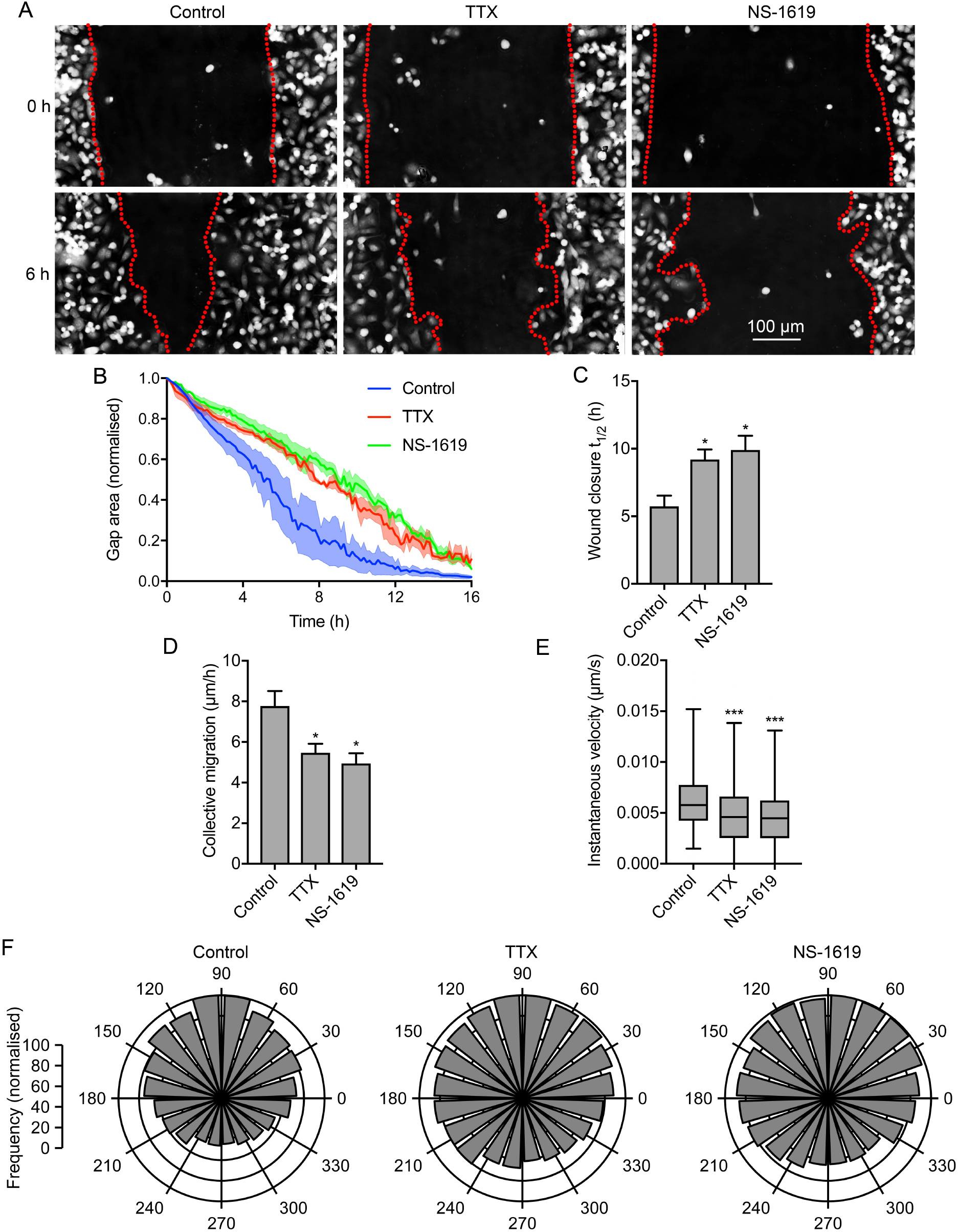
Na_v_1.5-dependent membrane potential depolarization regulates cell migration. (A) Representative scratch wounds at 0 h and 6 h into a wound healing assay ± TTX (30 μM) or NS-1619 (1 μM). Red dotted lines highlight wound edges. (B) Wound area during the migration assay (“gap remaining”), normalized to starting value (n = 3). (C) t_1/2_ of wound closure (n ≥ 5). (D) Collective migration (µm/h) of cells closing the wound (n ≥ 5). (E) Instantaneous velocity (µm/s) of segmented cells (n ≥ 2662). (F) Polar histograms showing directionality of migrating cells at the leading edge of wounds (P <0.001; Friedman with Dunn’s test). 90° = axis perpendicular to wound. Data in (B – D) are mean and SEM. Box plot whiskers in (E) show maximum and minimum values and horizontal lines show 75th, 50th, and 25th percentile values. *P < 0.05; **P < 0.01; ***P <0.001; ANOVA with Tukey test (C, D); Kruskal-Wallis with Dunn’s test (E).

Na_v_1.5 promotes an elongate, mesenchymal-like motile morphology (Brisson et al., 2013; Driffort et al., 2014; Nelson, Yang, Millican-Slater, et al., 2015). This morphological modulation has been shown to occur through two potentially overlapping mechanisms: (1) regulation of adhesion-mediated signaling *via* β1 subunits and up-regulation of CD44 expression (Nelson et al., 2014; Nelson, Yang, Millican-Slater, et al., 2015), and (2) *via* increased Src kinase activity, phosphorylation of the actin-nucleation-promoting factor cortactin, and F-actin polymerization (Brisson et al., 2013). In addition, the V_m_ has been shown to promote reorganization of the actin filament network and cytoskeleton in other cell types (Chifflet et al., 2004; Chifflet et al., 2005; Chifflet et al., 2003; Nin et al., 2009; Szaszi et al., 2005). However, the potential involvement of V_m_ in Na_v_1.5-mediated signaling is not known. We thus investigated the participation of Na_v_1.5-mediated V_m_ depolarization in regulating cell morphology. TTX (30 µM) significantly increased cell circularity after 3 h (P < 0.001; n ≥ 61; ANOVA with Tukey test; Figure 6A, B), consistent with previously published data (Brisson et al., 2013). NS-1619 (1 µM) also increased circularity (P < 0.05; n ≥ 61; ANOVA with Tukey test; Figure 6A, B). Similarly, TTX and NS-1619 both significantly reduced the Feret’s diameter (maximum caliper distance across the cell; P < 0.01; n ≥ 57; ANOVA with Tukey test; Figure 6C). These data suggest that V_m_ depolarization promotes acquisition of an elongate phenotype.

**Figure 6.**
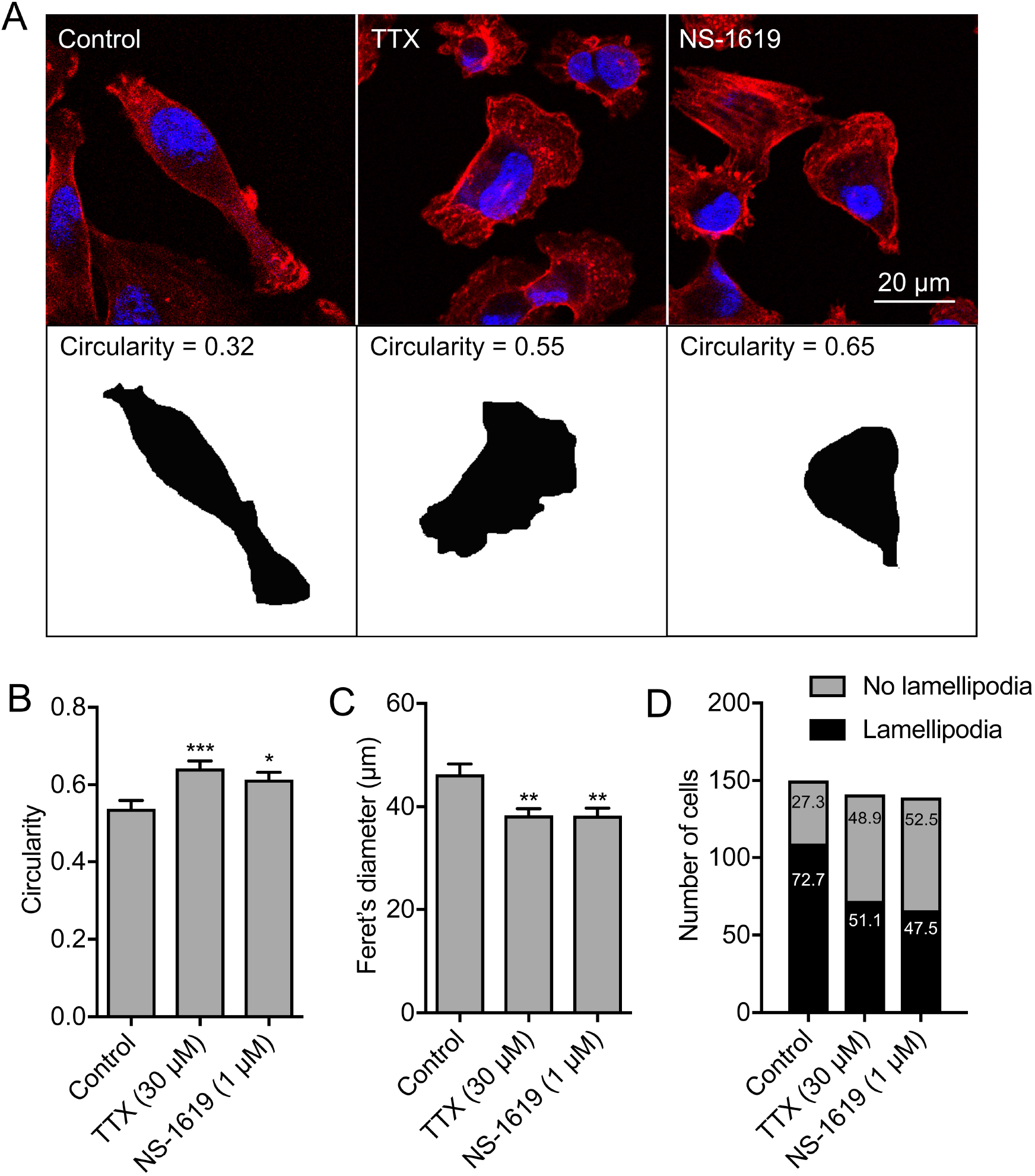
Na_v_1.5-dependent membrane potential depolarization regulates lamellipodia formation. (A) Images of representative cells after treatment with TTX (30 μM) or NS-1619 (1 μM) for 3 h. Cells were fixed and stained with phalloidin (red) and DAPI (blue). Lower row shows masks of cells in the upper row, from which the circularity was calculated. (B) Circularity (n ≥ 61). (C) Feret’s diameter (µm; n ≥ 57). (D) Number of MDA-MB-231 cells with a lamellipodium (P < 0.001; χ^2^ test). Numbers in bars are %. Bars in (B) and (C) are mean and SEM. *P < 0.05; **P < 0.01; ***P < 0.001; ANOVA with Tukey test.

Given that Na_v_1.5-mediated V_m_ depolarization promotes both morphological changes and cellular migration, we postulated that it might regulate formation of lamellipodia in migrating cells. We therefore scored the number of migrating cells with visible lamellipodia in a wound healing assay following treatment with TTX and NS-1619. Both treatments significantly reduced the proportion of cells with a lamellipodium in the measured population (P < 0.001; n ≥ 139 cells per condition; χ^2^ test; Figure 6D). Together, these findings suggest that Na_v_1.5-mediated V_m_ depolarization promotes a change in cellular morphology towards an elongate phenotype, and increases lamellipodia formation, thus potentiating cellular migration.

### Na_v_1.5-dependent membrane potential depolarization regulates Rac1 activation

The small GTPase Rac1, together with Rho and cdc42, plays a critical role in regulating cytoskeletal organization and cell motility (Burridge & Wennerberg, 2004). In addition, Rac1 also regulates the formation of lamellipodia, cell motility and the directionality of cell movement (Wu et al., 2009). We therefore postulated that Na_v_1.5 and V_m_ depolarization may regulate the level of active (GTP-bound) Rac1 in the lamellipodia of migrating cells. We treated migrating cells with NS-1619 and TTX for 3 h and then evaluated distribution of Rac1-GTP by quantifying immunocytochemical signal density across concentric arcs in the lamellipodium of individual cells (Figure 7A, B). As reported previously, a large quantity of Rac1 is present intracellularly in the perinuclear region, although there is also an enrichment of Rac1-GTP in the lamellipodium, consistent with its critical role in this region (Garcia-Mata et al., 2011; Wu et al., 2009). Both TTX and NS-1619 significantly reduced the peak Rac1-GTP signal at the leading edge of migrating cells (P < 0.05; n ≥ 66; 1-way ANOVA with Tukey test; Figure 7C). However, the level of total cellular Rac1-GTP, determined in whole cell lysates, was unaffected by TTX treatment (P = 0.80; n = 6; t test; Figure 7D). Given that the majority of Rac1 is present intracellularly (Figure 7A) (Das et al., 2015; Moissoglu, Slepchenko, Meller, Horwitz, & Schwartz, 2006), these data suggest that Na_v_1.5 may regulate Rac1 locally at the plasma membrane and/or within lamellipodia. We next evaluated the effect of TTX and NS-1619 on distribution of Rac1-GTP *vs.* total Rac1 using an antibody that does not distinguish between GDP- and GTP-bound forms of Rac1 (Figure 7E). The peak total Rac1 signal at the leading edge of migrating cells was unaffected by both TTX and NS-1619 (P = 0.69; n ≥ 60; 1-way ANOVA; Figure 7F, G). Thus, the ratio of peak Rac1-GTP to total Rac1 at the leading edge was significantly reduced by both TTX and NS-1619 (P < 0.001; n = 3; 1-way ANOVA; Figure 7H). These data suggest that V_m_ depolarization caused by steady state Na_v_1.5 activity increases Rac1 activation at the leading edge of migrating cells.

**Figure 7.**
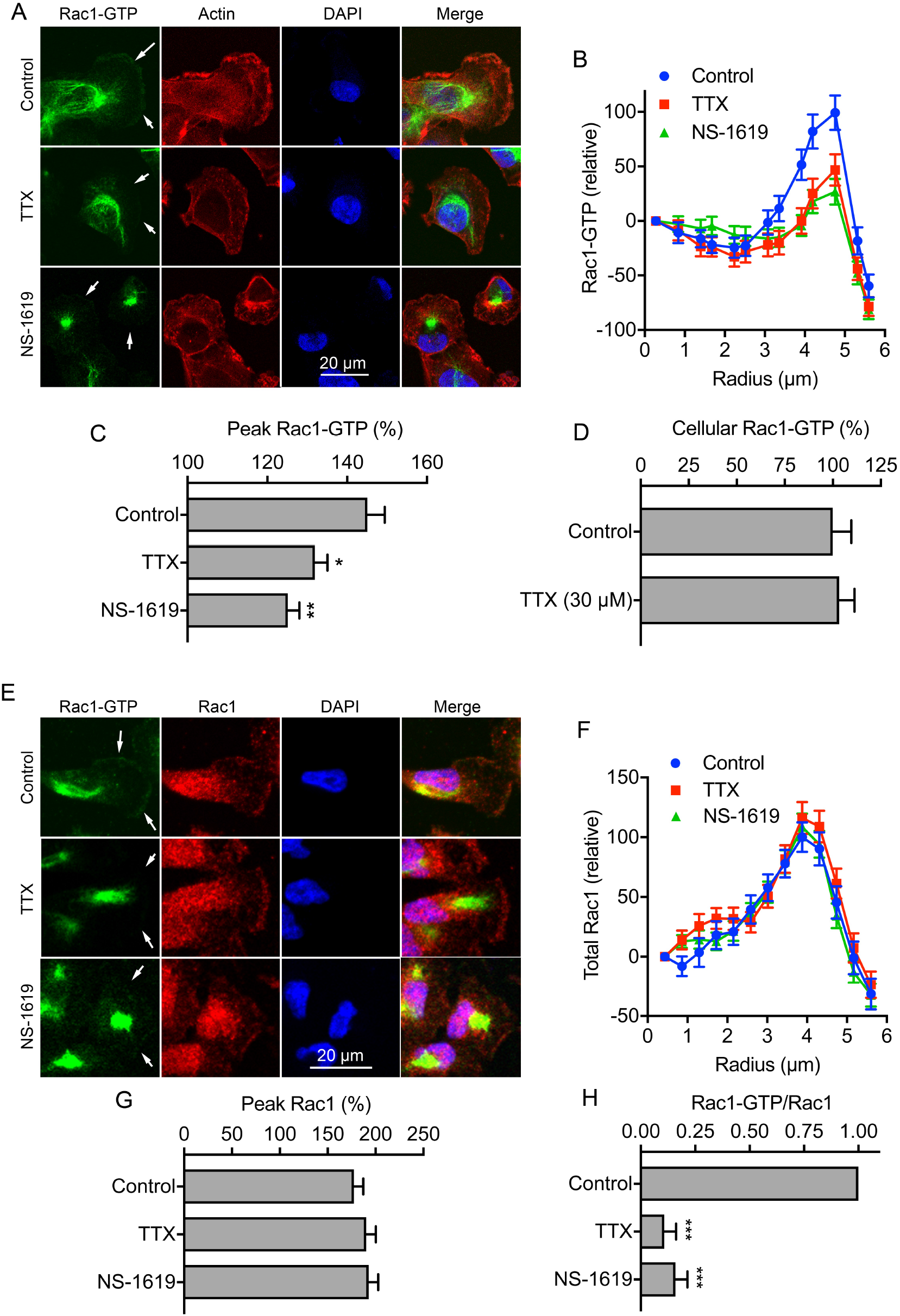
Na_v_1.5 and V_m_ regulate Rac1 activation/distribution. (A) Images of representative cells after treatment with TTX (30 µM) and NS-1619 (1 µM) for 3 h. Cells were labeled with Rac1-GTP antibody (green), phalloidin (red) and DAPI (blue). Arrows in the Rac1-GTP panels highlight the distribution or lack of expression at the leading edge. (B) Rac1-GTP signal density, measured across 20 arcs, in 0.43 µm radius increments, within a quadrant mask region of interest at the leading edge, normalized to the first arc (n ≥ 66). (C) Peak Rac1-GTP signal density per cell from (B), normalized to the first arc (n ≥ 66). (D) Total Rac1-GTP quantified in whole cell lysates using colorimetric small GTPase activation assay (n = 6). (E) Images of representative cells after treatment with TTX (30 µM) and NS-1619 (1 µM) for 3 h. Cells were labeled with Rac1-GTP antibody (green), total Rac1 antibody (red), and DAPI (blue). Arrows in the Rac1-GTP panels highlight the distribution or lack of expression at the leading edge. (F) Total Rac1 signal density, measured across 20 arcs, in 0.43 µm radius increments, within a quadrant mask region of interest at the leading edge, normalized to the first arc (n ≥ 59). (G) Peak Rac1 signal density per cell from (F), normalized to the first arc (n ≥ 59). (H) Ratio of Peak Rac1-GTP/Peak total Rac1 for each experimental repeat, normalized to control (n = 3). Data are mean and SEM. *P < 0.05; **P < 0.01; ***P < 0.001; ANOVA with Tukey test.

V_m_ depolarization promotes clustering and activation of the small GTPase K-Ras as a result of voltage-dependent redistribution of charged phospholipids including phosphatidylserine (Zhou et al., 2015). Given that Rac1 is also anchored to the inner leaflet of the plasma membrane via interaction with phosphatidylserine (Finkielstein et al., 2006; Magalhaes & Glogauer, 2010), we reasoned that a similar voltage-dependent mechanism may be responsible for Rac1 activation in the lamellipodia. We therefore next evaluated the effect of V_m_ depolarization on colocalization of Rac1-GTP with phosphatidylserine, using annexin V as a marker (Figure 8A). Treatment of migrating cells with TTX and NS-1619 reduced colocalization of Rac1-GTP with phosphatidylserine in the lamellipodia (Figure 8B-D). This was confirmed by quantification of thresholded Manders colocalization coefficients for phosphatidylserine and Rac1-GTP (P < 0.01; n = 30; 2-way ANOVA with Tukey test; Figure 8E). Similarly, the TTX and NS-1619 treatments both reduced the Li’s intensity correlation quotient (P < 0.001; n = 30; 1-way ANOVA with Tukey test; Figure 8F). These data support the notion that V_m_ depolarization increases Rac1 localization in the lamellipodia as a result of its interaction with phosphatidylserine.

**Figure 8.**
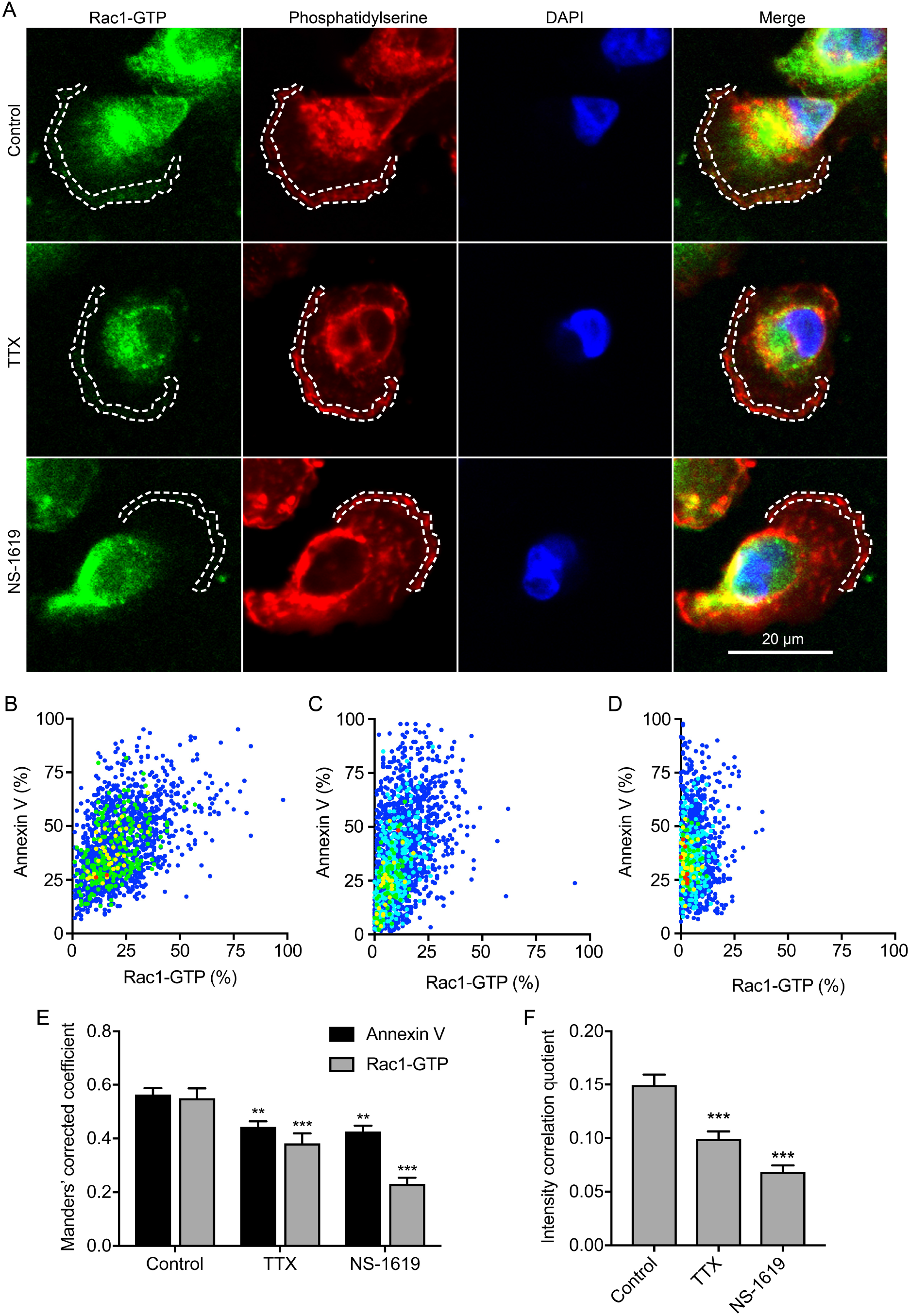
Na_v_1.5 and V_m_ regulate Rac1-GTP colocalization with phosphatidylserine. (A) Images of representative cells after treatment with TTX (30 µM) and NS-1619 (1 µM) for 3 h. Cells were labeled with Rac1-GTP antibody (green), annexin V (red) and DAPI (blue). Dashed lines highlight regions of interest at the leading edge. (B) Cytofluorogram showing colocalization of annexin V and Rac1-GTP staining in region of interest in control cell from (A), normalized to maximum in each channel. (C) Cytofluorogram showing colocalization of annexin V and Rac1-GTP staining in region of interest in TTX cell from (A), normalized to maximum in each channel. (D) Cytofluorogram showing colocalization of annexin V and Rac1-GTP staining in region of interest in NS-1619 cell from (A), normalized to maximum in each channel. (E) Manders’ corrected colocalization coefficients for annexin V and Rac1-GTP staining in regions of interest of cells after treatment with TTX (30 µM) and NS-1619 (1 µM) for 3 h (n = 30). (F) Li’s intensity correlation quotient for Rac1-GTP and annexin V colocalization (n = 30). Data are mean and SEM. **P < 0.01; ***P < 0.001; ANOVA with Tukey test.

In order to evaluate whether Na_v_1.5-mediated V_m_ depolarization promotes Rac1 activity in live cells, and to independently confirm the immunocytochemistry data, we next employed a genetically encoded Rac1 FRET biosensor to monitor Rac1 activation. In agreement with a previous report (Fritz et al., 2015), the biosensor showed a general gradient of Rac1 activation increasing from the cell interior towards the periphery, consistent with its critical role in actin remodeling at the edge of migrating cells (Figure 9A) (Wu et al., 2009). Treatment with TTX did not affect biosensor distribution (Figure 9A), but suppressed activation at the periphery (Figure 9B, C). Quantification of FRET across the whole cell revealed that TTX significantly reduced Rac1 activation to 48.6 ± 5.2 % of control (P < 0.001; n ≥ 49; t test; Figure 9D). Together, these results indicate that blockade of Na_v_1.5 channels suppresses Rac1 activation at the cell periphery.

**Figure 9.**
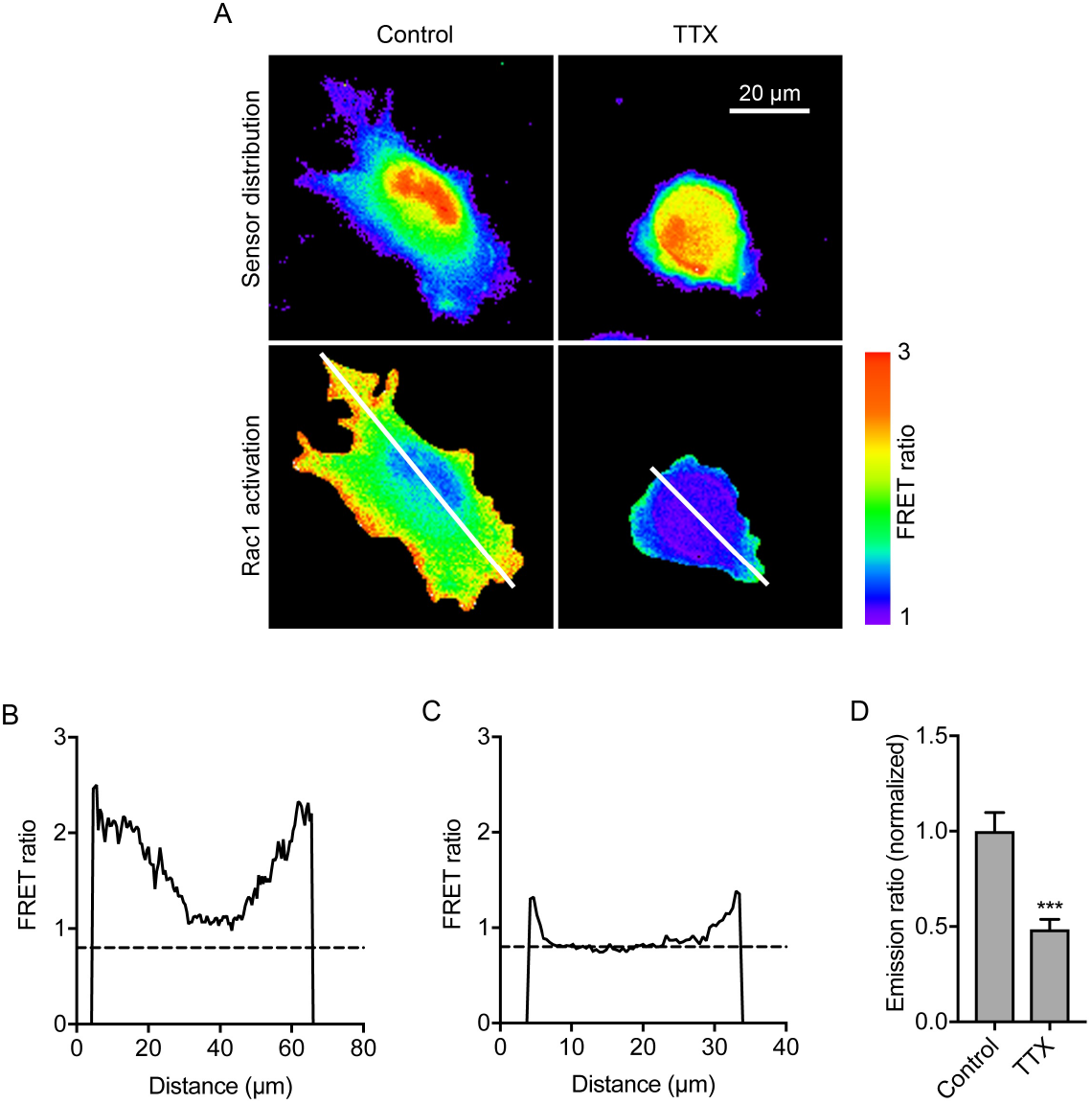
Na_v_1.5 regulates Rac1 activation in live cells detected using a genetically encoded Rac1 FRET biosensor. (A) Images of representative cells after treatment ± TTX (30 µM) for 3 h. Rac1 activation biosensor distribution is shown in the donor (mTFP) channel. Images are color-coded so that warm and cold colors represent high and low values, respectively, for sensor distribution and activation (FRET). (B) Fluorescence intensity profile along line drawn across control cell in (A). (C) Fluorescence intensity profile along line drawn across TTX-treated cell in (A). (D) Emission ratio (FRET), for cells measured after treatment ± TTX (30 µM) for 3 h, normalized to control (n ≥ 33). Data are mean and SEM. **P < 0.01; Student’s t test.

### Na_v_1.5-mediated morphological changes are dependent on Rac1 activation

Given that Na_v_1.5-dependent V_m_ depolarization promotes morphological changes leading to increased cell migration (Figures 5, 6), and that Na_v_1.5 promotes activation of Rac1 (Figures 7, 8, 9), which in turn, plays a key role in regulating cellular morphology and migration (Wu et al., 2009), we analyzed the impact of Rac1 inhibition on Na_v_1.5-dependent changes in cell morphology (Figure 10A). The specific Rac1 inhibitor EHT1864 increased circularity and reduced Feret’s diameter in a dose-dependent manner (Figure S5A, B). Furthermore, EHT1864 increased circularity to a similar extent to TTX (P < 0.001; n ≥ 308; 1-way ANOVA with Tukey test; Figure 10B, C). Importantly, co-application of EHT1864 with TTX had no additive effect on circularity or Feret’s diameter (Figure 10B, C). These results show that Rac1 activity is required to transduce the Na_v_1.5-dependent V_m_ depolarizing signal in order to promote acquisition of an elongate, motile cell phenotype (Figure 10D).

**Figure 10.**
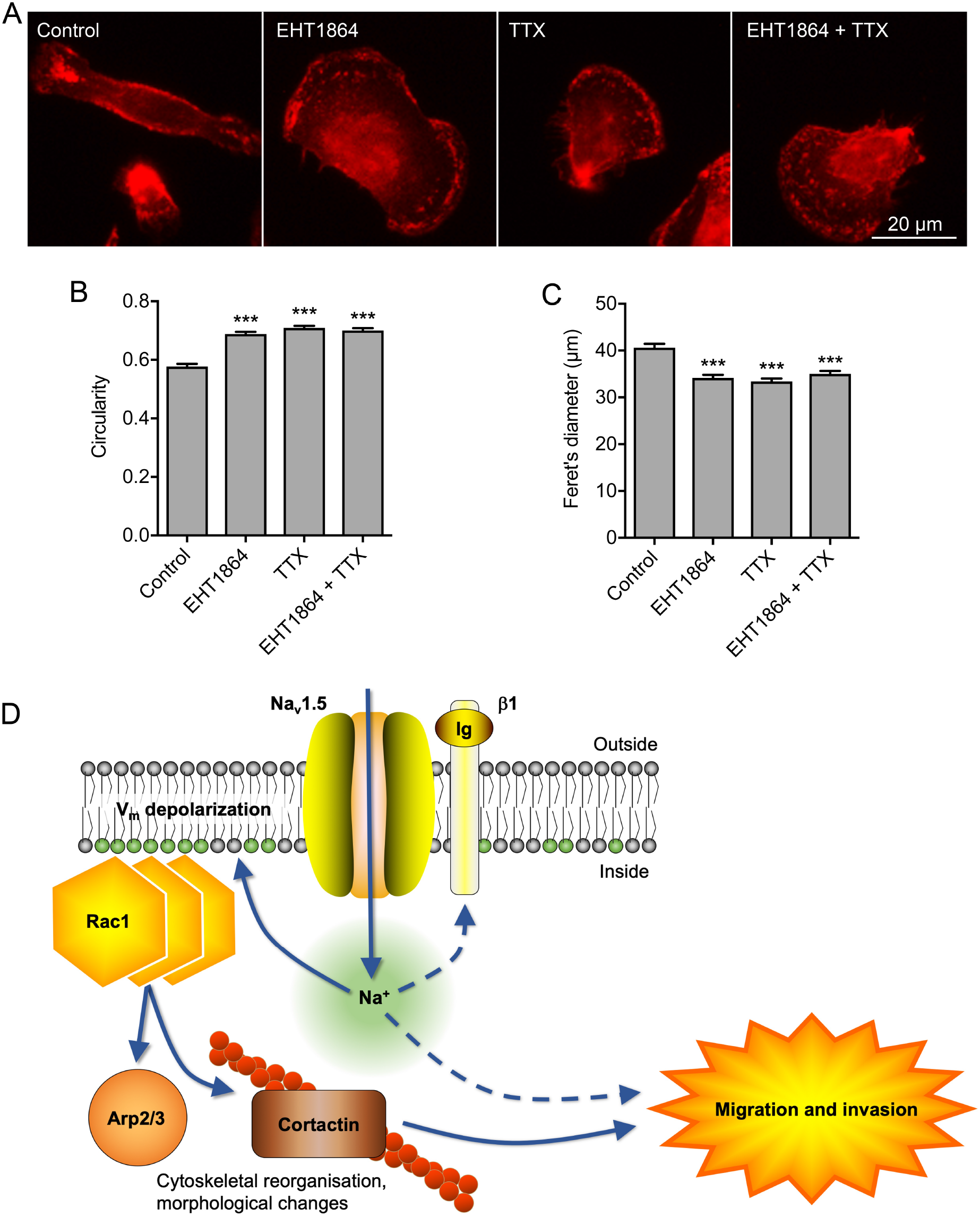
Na_v_1.5-mediated morphological changes and migration are dependent on Rac1 activation. (A) Images of representative cells after treatment with TTX (30 µM) ± Rac1 inhibitor (EHT1864; 0.5 µM) for 3 h. Cells are labeled with CD44 antibody (red). (B) Circularity of cells after treatment with TTX (30 µM) ± EHT1864 (0.5 µM) for 3 h (n ≥ 308). (C) Feret’s diameter (µm) of cells after treatment with TTX (30 µM) ± EHT1864 (0.5 µM) for 3 h (n ≥ 308). Data are mean and SEM. ***P < 0.001; ANOVA with Tukey test. (D) Proposed mechanism underlying Na_v_1.5-mediated V_m_-dependent morphological changes and migration. Na_v_1.5 channels carry Na^+^ influx, which depolarizes the V_m_, causing redistribution of charged phosphatidylserine in the inner leaflet of the phospholipid bilayer, promoting Rac1 redistribution and activation. Rac1 regulates cytoskeletal modification *via* the Arp2/3 complex and increasing phosphorylation of cortactin and cofilin, promoting acquisition of a promigratory phenotype (Pollard, 2007; Stock & Schwab, 2015). Na^+^ influx through Na_v_1.5 channels may also impact on migration and invasion through other mechanism(s), including via β1 subunit-dependent adhesion (Brisson et al., 2013; Carrie D House et al., 2010; C. D. House et al., 2015; Nelson et al., 2014).

## Discussion

Here we identify Na_v_1.5-mediated V_m_ depolarization as a regulator of Rac1 activation. Thus, we link ionic and electrical signaling at the plasma membrane to small GTPase-dependent cytoskeletal reorganization and cellular migration. We show that depletion of extracellular Na^+^ or blockage of Na_v_1.5 channels reversibly hyperpolarizes the V_m_ at the timescale of minutes. We further show that Na_v_1.5-dependent V_m_ depolarization increases Rac1 colocalization with phosphatidylserine at the leading edge of migrating cells, promoting Rac1 activation, and resulting in acquisition of a motile, mesenchymal-like cellular phenotype. We therefore propose that Na_v_1.5 may serve as a sensor for local changes in the ionic microenvironment, thus permitting voltage-dependent activation of Rac1 to fine tune cell migration.

### Ion conductance, membrane potential and migration

We show that Na_v_1.5 carries a persistent inward Na^+^ current in MDA-MB-231 cells. In agreement with the work of others, we report incomplete inactivation of Na_v_1.5 resulting in a small voltage window encompassing the V_m_, thus supporting this persistent Na^+^ current (Brisson et al., 2011; Djamgoz & Onkal, 2013; Gillet et al., 2009; Nelson, Yang, Millican-Slater, et al., 2015; Roger et al., 2003; Yang et al., 2012). The persistent inward Na^+^ current carried by Na_v_1.5 contributes to steady-state V_m_ depolarization. Our findings agree with another study in H460 non-small cell lung cancer cells, in which Na_v_1.7 channels were shown to depolarize the V_m_ by ∼10 mV (Campbell et al., 2013). Thus, the mechanism identified here is likely applicable to other non-excitable cell types where VGSCs are expressed (Chen et al., 2019; Grimes et al., 1995; Carrie D House et al., 2010; Persson et al., 2014; Zhou et al., 2015). In addition, it has previously been shown that VGSCs contribute to steady-state V_m_ depolarization and viability in rat optic nerve axons and spinal cord astrocytes over a time course of hours (Sontheimer, Fernandez-Marques, Ullrich, Pappas, & Waxman, 1994; Stys, Sontheimer, Ransom, & Waxman, 1993), suggesting that this mechanism persists over an extended period. There may be other Na^+^-permeable pathway(s) also contributing to V_m_ regulation in non-excitable cells, e.g. via epithelial Na^+^ channels (Amara, Ivy, Myles, & Tiriveedhi, 2015), Na^+^-K^+^ ATPase (Winnicka et al., 2008), and NHE1 (Brisson et al., 2011). The V_m_ may also be dependent on the activity of transporters and channels regulating the movement of other ions, including Cl^-^ and K^+^ (Yang & Brackenbury, 2013). The depolarized resting V_m_ reported here may therefore be attributed not only to the high permeability to Na^+^ but also to the low expression/activity of hyperpolarizing K^+^ channels.

Relatively small alterations in V_m_ can be functionally significant. For example, during synaptic transfer from photoreceptors to ganglion cells in turtle retina, V_m_ depolarizations as small as 5 mV are sufficient to activate paired ganglion cells (Baylor & Fettiplace, 1977). The V_m_ can function as an instructive signal to regulate cell cycle progression in proliferating cells (Cervera et al., 2016; Cone, 1971; Cone & Cone, 1976; Zhou et al., 2015). V_m_ depolarization can regulate other cellular behaviors, including differentiation (Sundelacruz et al., 2008) and cytoskeletal reorganization (Chifflet et al., 2004; Chifflet et al., 2005; Chifflet et al., 2003; Nin et al., 2009; Szaszi et al., 2005). At the tissue level, changes in V_m_ can also regulate morphogenesis, regeneration and tumorigenesis (Beane et al., 2011; Beane et al., 2013; Cervera et al., 2016; Chernet et al., 2016; Chernet & Levin, 2014; Lobikin et al., 2012; Lobikin et al., 2015). Several studies have shown that epithelial cells undergo V_m_ depolarization when migrating into wounds (Chifflet et al., 2004; Chifflet et al., 2005). However, an interesting and novel finding of our study here is that V_m_ depolarization is not simply a consequence of motile behavior (Schwab et al., 2012), but is *itself* a master regulator of morphological changes and migration. Here, we found that hyperpolarizing the V_m_ by ∼5 mV reduced migration by ∼30 %. The fact that this level of V_m_ hyperpolarization did not completely inhibit migration raises the possibility that additional hyperpolarization may have a further inhibitory effect. It is also likely that other ion conductance routes contributing to, and/or dependent on the V_m_, may be involved in migration regulation. For example, hyperpolarization would increase the driving force for Ca^2+^ entry. Nonetheless, our data underscore that persistent Na^+^ influx through Na_v_1.5 channels leads to V_m_ depolarization, which in turn promotes cell migration.

Given the general trend towards depolarized V_m_ both in cancer cells (Cervera et al., 2016; Fraser et al., 2005; Yang & Brackenbury, 2013), and in epithelial cells undergoing migration (Chifflet et al., 2004; Chifflet et al., 2005), we propose that voltage-dependent migratory behavior may be a general cellular phenomenon. For example, although involvement of V_m_ was not directly investigated, activation of the V_m_-hyperpolarizing channel K_Ca_1.1 with NS-1619 reduces migration of glioma cells (Kraft et al., 2003). Furthermore, VGSCs have been detected in cells from a broad range of tumor types, where they potentiate migration and invasion (Besson et al., 2015; Brackenbury, 2012). We therefore argue that the results presented here have broad applicability to other cell types.

### Voltage-dependent regulation of Rac1 activation

V_m_ depolarization promotes K-Ras clustering and subsequent activation (Zhou et al., 2015). Various small GTPases, including K-Ras and the Rho GTPases, are anchored to the inner leaflet of the plasma membrane through interaction with the anionic phospholipids PIP2, PIP3 and phosphatidylserine (Finkielstein et al., 2006; Heo et al., 2006; Magalhaes & Glogauer, 2010; Remorino et al., 2017; Zhou et al., 2015). In the case of K-Ras, clustering and activation has been shown using electron microscopy to arise as a result of depolarization-induced nanoscale redistribution of phosphatidylserine and PIP2 in the inner leaflet of the phospholipid bilayer, to which the K-Ras is anchored (Zhou et al., 2015). Our major discovery here is that Rac1 activation is also voltage-dependent. Our data show that Na_v_1.5-mediated V_m_ depolarization causes Rac1 activation, likely as a result of voltage-dependent redistribution of phosphatidylserine (Figure 10D). Thus, V_m_-dependent signaling is not exclusively limited to mitogenic cascades but is also a key regulator of morphological changes and migration.

A growing body of evidence implicates V_m_ depolarization in regulation of small GTPase activity. For example, V_m_ has been shown to regulate the activity of GEF-H1, which, in turn, regulates the activity of the Rho/Rho kinase pathway (Waheed et al., 2010). V_m_ depolarization-dependent activation of Rho and subsequent cytoskeletal organization has been shown to be Ca^2+^-independent (Szaszi et al., 2005). In agreement with this, our data also imply that Ca^2+^ is not involved as a signaling intermediary in V_m_ depolarization-induced Rac1-mediated promotion of migration. On the other hand, in ATP-stimulated microglia, VGSC activity increases [Ca^2+^]_i_, and activates ERK and Rac1, promoting migration (Persson et al., 2014). V_m_ depolarization-induced activation of Ras and Rap1 has also been reported in mouse cortical neurons (Baldassa, Zippel, & Sturani, 2003).

The mechanism uncovered here may feed into, and interact with, additional signaling pathways which have been shown to be dependent on Na_v_1.5 activity. For example, Na_v_1.5 has been shown to promote Src kinase activity and phosphorylation of cortactin and cofilin (Brisson et al., 2013). Thus, Na_v_1.5-induced V_m_ depolarization may activate Rac1, increasing cortactin phosphorylation and therefore enhancing cofilin activity and actin filament polymerization. Src has also been shown to regulate Rac1 activity, suggesting the potential for feedback regulation (Servitja, Marinissen, Sodhi, Bustelo, & Gutkind, 2003). Na_v_1.5 positively regulates the expression of the metastasis-promoting protein CD44, which may activate Src and therefore contribute to this process (Bourguignon, Zhu, Shao, & Chen, 2001; Carrie D House et al., 2010; C. D. House et al., 2015; Lee, Wang, Sudhir, & Chen, 2008; Nelson, Yang, Millican-Slater, et al., 2015). In addition, Na_v_1.5 is expressed on astrocytes where it regulates migration by promoting reverse Na^+^/Ca^2+^ exchange (J. A. Black, Dib-Hajj, Cohen, Hinson, & Waxman, 1998; Pappalardo, Samad, Black, & Waxman, 2014). Furthermore, Na_v_1.5 is expressed on the endosome of macrophages, where it regulates endosomal acidification (Carrithers et al., 2007). On the other hand, Na_v_1.6 is expressed within macrophage podosomes, where it promotes invasion via intracellular Na^+^ release (Carrithers et al., 2009). Na_v_1.5 and Na_v_1.6 are also expressed on microglia, where the latter has been shown to also regulate migration and cytokine release (Joel A. Black, Liu, & Waxman, 2009). It remains unknown whether Na_v_1.5 opening changes during migration and what the signals might be that would regulate such alterations; however, it has been shown that Na_v_1.5 can be regulated by growth factors and hormone signaling, and is itself mechanosensitive (Beyder et al., 2010; Fraser et al., 2014). Together, these studies suggest that VGSCs may further regulate the behavior of non-excitable cells via pathways in addition to the one identified here.

### Implications for our understanding of metastasis

We provide evidence that Na_v_1.5 contributes to steady-state V_m_ depolarization, which in turn promotes V_m_-dependent Rac1 activation, lamellipodial protrusion formation and cellular migration. These data are in agreement with the previously reported role for VGSCs in regulating morphology and migration of tumor cells (Brackenbury et al., 2007; Brackenbury & Djamgoz, 2006; Brisson et al., 2013; Fraser, Ding, Liu, Foster, & Djamgoz, 1999; Fraser et al., 2005; Nelson, Yang, Dowle, et al., 2015). It is therefore conceivable that depolarization-dependent Rac1 activation may contribute to metastatic dissemination in response to local ionic changes in the tumor microenvironment. Thus, pharmacological targeting of VGSCs, which inhibits metastasis in preclinical models (Driffort et al., 2014; Nelson, Yang, Dowle, et al., 2015; Yildirim, Altun, Gumushan, Patel, & Djamgoz, 2012), may provide therapeutic benefit via V_m_ hyperpolarization and downregulation of V_m_-dependent small GTPase activation. In conclusion, our results reveal a new role for Na_v_1.5 channels as voltage-dependent activators of Rac1 signaling to promote cellular migration.

## Author contributions

MY and WB contributed to the conception and design of the work. MY, AJ, RK, RS, PO’T and WB contributed to acquisition, analysis, and interpretation of data for the work. MY, RK, RS, PO’T and WB contributed to drafting the work and revising it critically for important intellectual content. All authors approved the final version of the manuscript.

## Acknowledgements

This work was supported by the Medical Research Council [G1000508], Cancer Research UK [A25922] and a Radhika V Sreedhar Scholarship.

