## Supplementary Figures for "Voltage-dependent activation of Rac1 by Na_v_1.5 channels promotes cell migration"

### Supplementary Figure Legends

**Figure S1.** Effects of sodium citrate and phenytoin on the membrane potential. (A) $V_m$ in control physiological saline solution, after perfusion with the vehicle for TTX (148 μM sodium citrate, pH = 4.8) and following washout. Solid line, mean; gray shading, SEM (n = 13). (B) Quantification of $V_m$ over the last 5 s in control, sodium citrate, and washout (n = 13). (C) Representative trace showing the inhibitory effect of phenytoin (100 μM) on $Na^+$ current, and recovery after washout. The cell was held at -120 mV for 250 ms before depolarizing to -10 mV for 50 ms. (D) Expanded view of persistent $Na^+$ current 40-45 ms following onset of depolarization. (E) Quantification of the normalized transient $Na^+$ current elicited by depolarizing to -10 mV from a holding potential of -120 mV (n = 9). (F) Quantification of the normalized persistent $Na^+$ current 40-45 ms after depolarizing to -10 mV from a holding potential of -120 mV (n = 4). (G) Quantification of the normalized transient $Na^+$ current elicited by depolarizing to -10 mV from a holding potential of -80 mV (n = 4). (H) Quantification of the normalized persistent $Na^+$ current 40-45 ms after depolarizing to -10 mV from a holding potential of -80 mV (n = 3). (I) $V_m$ in control physiological saline solution, after phenytoin (100 μM) treatment and following washout. Solid line, mean; gray shading, SEM (n = 12). (J) Quantification of $V_m$ over the last 5 s in control, phenytoin, and washout (n = 12). (K) Quantification of $V_m$ over the last 5 s in control, NaOH (75 μM; vehicle for phenytoin) and washout (n = 9). Data are mean and SEM. *P < 0.05; **P < 0.01; ***P < 0.001; repeated measures ANOVA with Tukey test.

**Figure S2.** Effects of veratridine on $Na^+$ current and NMDG on membrane potential. (A) I-V relationship for transient $Na^+$ current in control physiological saline solution and following perfusion of veratridine (100 μM) (n = 6). (B) I-V relationship for persistent $Na^+$ current (defined as mean current density 45-50 ms following onset of depolarization; n = 6). Data are mean and SEM. Consistent with previous reports [1,2], veratridine caused a small reduction in the transient peak $Na^+$ current density but increased the peak persistent $Na^+$

current density. (C) $V_m$ in control physiological saline solution, after extracellular $Na^+$ replacement with N-methyl-D-glucamine (NMDG) and following washout. Solid line, mean; gray shading, SEM (n = 7). (D) Quantification of $V_m$ over the last 5 s in control, NMDG, and washout (n = 7).

**Figure S3.** Tetrodotoxin and NS-1619 do not affect the intracellular $Ca^{2+}$ level. (A) Intracellular $Ca^{2+}$ level (340/380 ratio) following perfusion with TTX (30 μM). Solid line, mean; gray shading, SEM (n = 55). (B) 340/380 ratio over the last 30 s in control, TTX and washout (n = 3). (C) Intracellular $Ca^{2+}$ level (340/380 ratio) following pre-treatment with TTX (30 μM) for 48 h (n = 55). (D) 340/380 ratio over the last 30 s of TTX and washout (n = 3). (E) Intracellular $Ca^{2+}$ level (340/380 ratio) following perfusion with NS-1619 (1 μM). Solid line, mean; gray shading, SEM (n = 40). (F) 340/380 ratio over the last 30 s in control, NS-1619 and washout (n = 3). Ionomycin was used at the end of all experiments as a positive control confirming sensitivity of the $Ca^{2+}$ indicator. Data are mean and SEM.

**Figure S4.** Effect of tetrodotoxin and NS-1619 on proliferation and invasion. (A) Proliferation (quantified as number of cells using the MTT assay) following treatment for 24 h with NS-1619 (1 μM, 40 μM) or vehicle (n = 3). (B) Matrigel invasion ± NS-1619 (1 μM) or TTX (30 μM), normalized to control (n = 4). (C) Matrigel invasion ± NS-1619 (40 μM), normalized to control (n = 3). Data are mean and SEM. *P < 0.05; **P < 0.01; repeated measures ANOVA with Tukey test.

**Figure S5.** Dose-dependent effect of EHT1864 on cell morphology. (A) Circularity of cells after treatment with EHT1864 (0.5-10 μM) or vehicle for 3 h (n ≥ 277). (C) Feret's diameter (μm) of cells after treatment with EHT1864 (0.5-10 μM) or vehicle for 3 h (n ≥ 277). Data are mean and SEM.

Figure S1

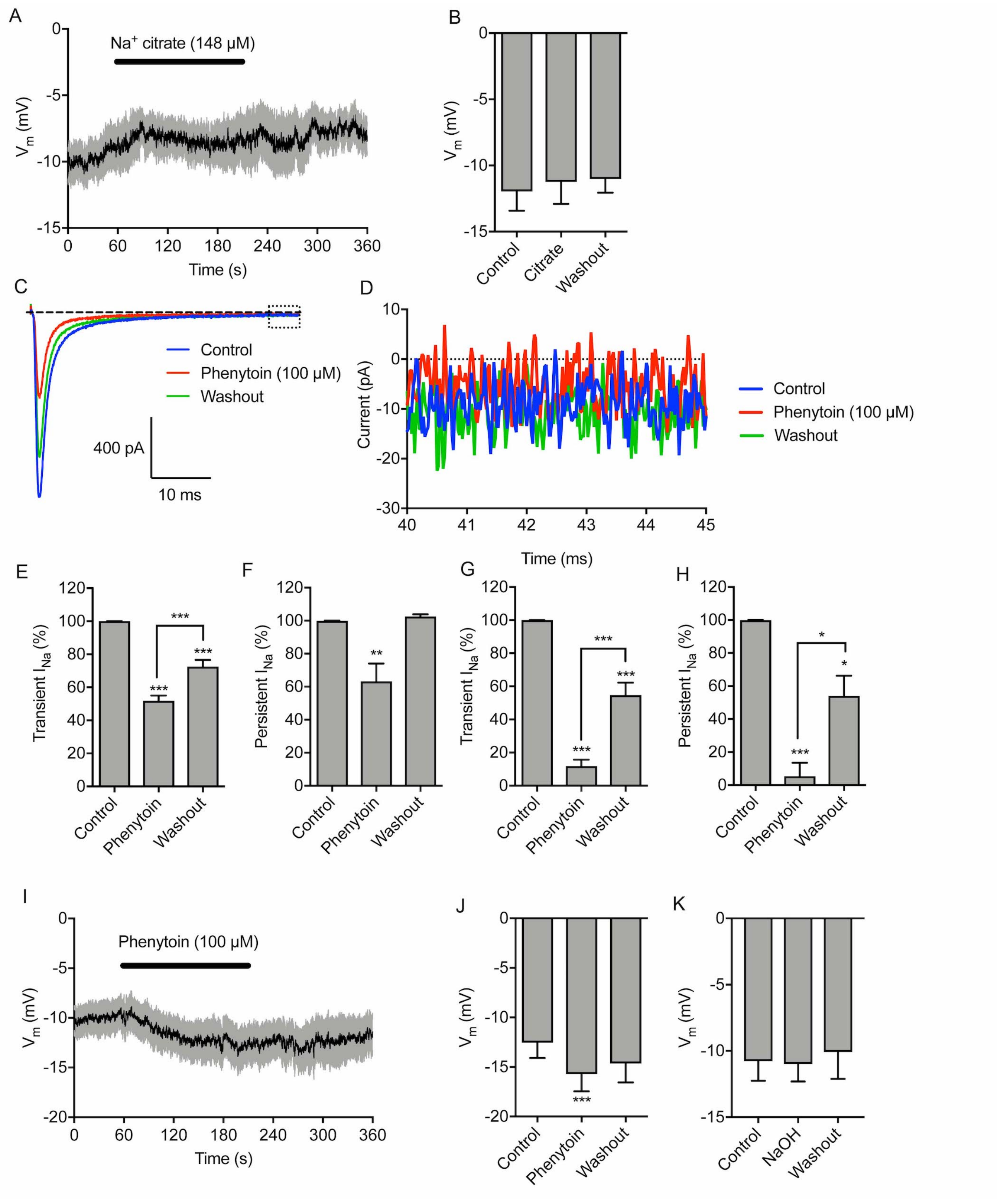

A
Na+ citrate (148 µM)
Vm (mV)
Time (s)
B
Vm (mV)
Control
Citrate
Washout
C
Control
Phenytoin (100 µM)
Washout
400 pA
10 ms
D
Current (pA)
Time (ms)
Control
Phenytoin (100 µM)
Washout
E
Transient INa (%)
Control
Phenytoin
Washout
F
Persistent INa (%)
G
Transient INa (%)
H
Persistent INa (%)
I
Phenytoin (100 µM)
Vm (mV)
Time (s)
J
Vm (mV)
Control
Phenytoin
Washout
K
Vm (mV)
Control
NaOH
Washout

Figure S2

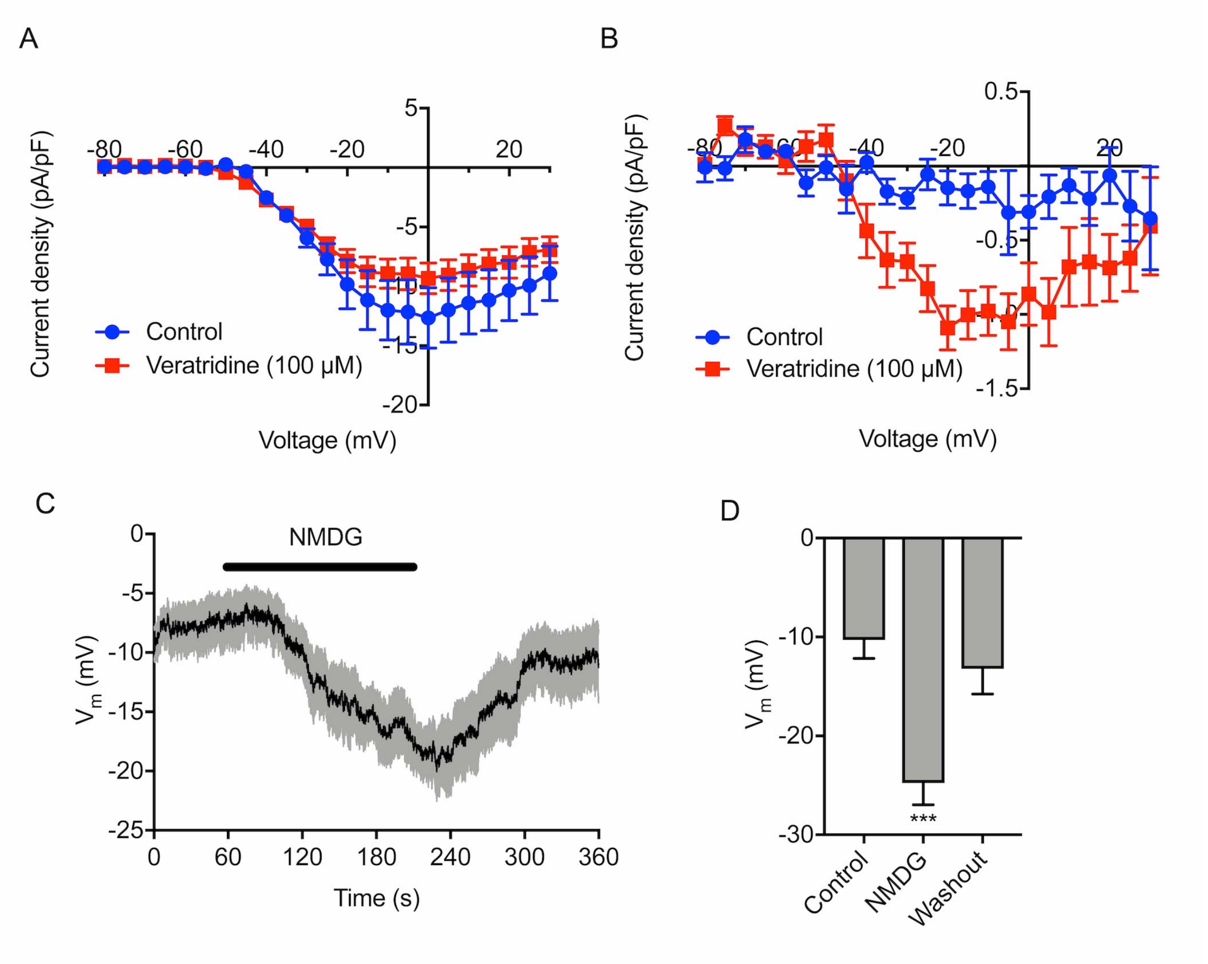
A
Current density (pA/pF)
Voltage (mV)
Control
Veratridine (100 µM)
B
Current density (pA/pF)
Voltage (mV)
Control
Veratridine (100 µM)
C
NMDG
$V_m$ (mV)
Time (s)
D
$V_m$ (mV)
***
Control
NMDG
Washout

Figure S3

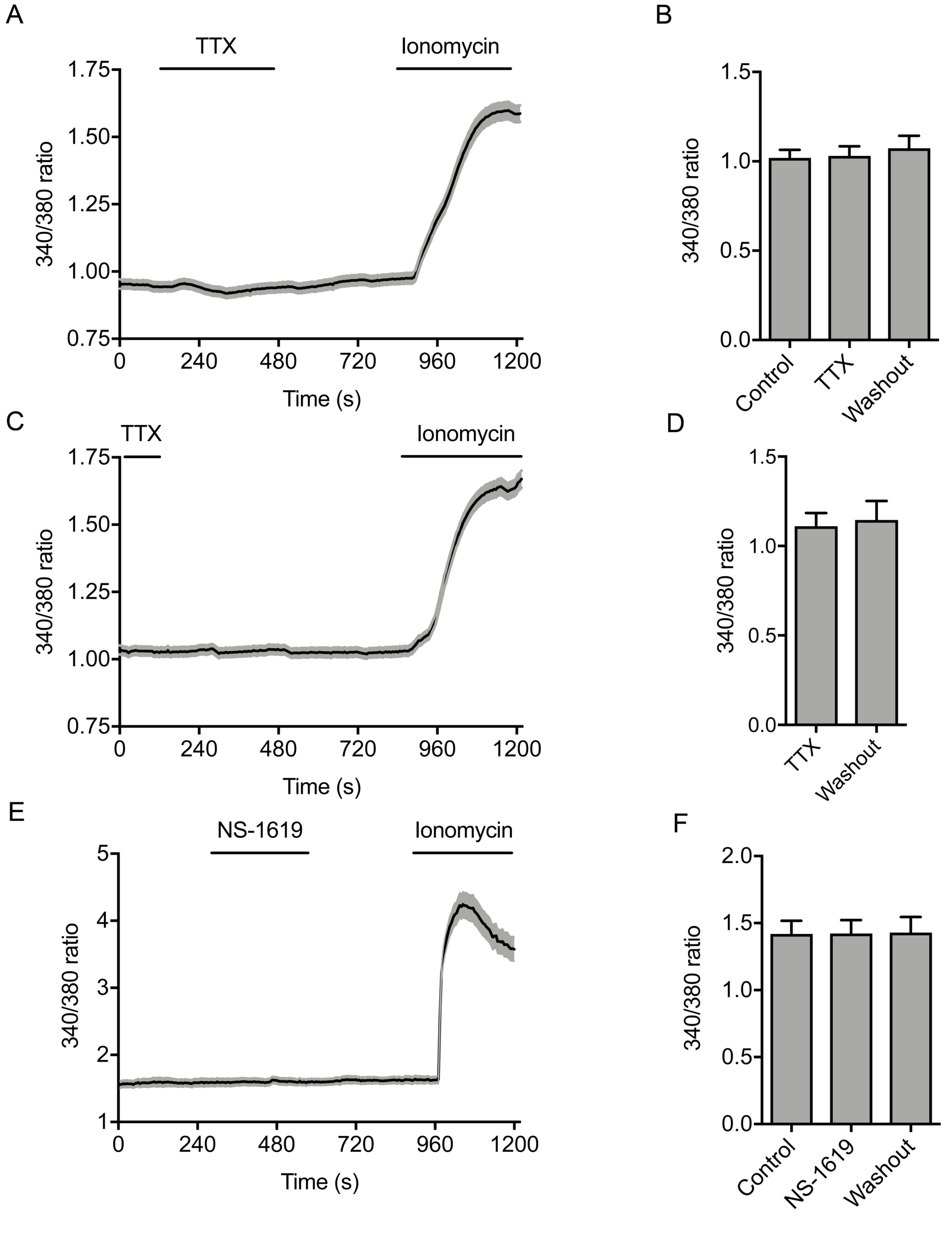
A
TTX
Ionomycin
1.75
1.50
1.25
1.00
0.75
340/380 ratio
0
240
480
720
960
1200
Time (s)
B
1.5
1.0
0.5
0.0
340/380 ratio
Control
TTX
Washout
C
TTX
Ionomycin
1.75
1.50
1.25
1.00
0.75
340/380 ratio
0
240
480
720
960
1200
Time (s)
D
1.5
1.0
0.5
0.0
340/380 ratio
TTX
Washout
E
NS-1619
Ionomycin
5
4
3
2
1
340/380 ratio
0
240
480
720
960
1200
Time (s)
F
2.0
1.5
1.0
0.5
0.0
340/380 ratio
Control
NS-1619
Washout

Figure S4

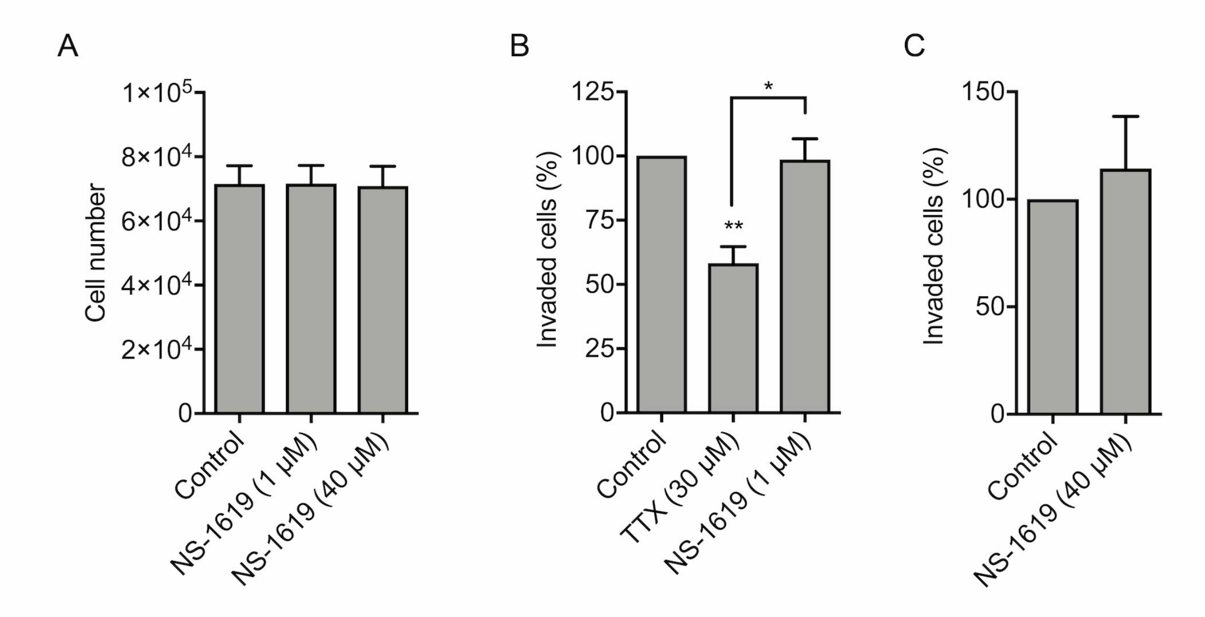

A
B
C
Cell number
1×10^5
8×10^4
6×10^4
4×10^4
2×10^4
0
Control
NS-1619 (1 μM)
NS-1619 (40 μM)
Invaded cells (%)
125
100
75
50
25
0
**
*
Control
TTX (30 μM)
NS-1619 (1 μM)
Invaded cells (%)
150
100
50
0
Control
NS-1619 (40 μM)

Figure S5

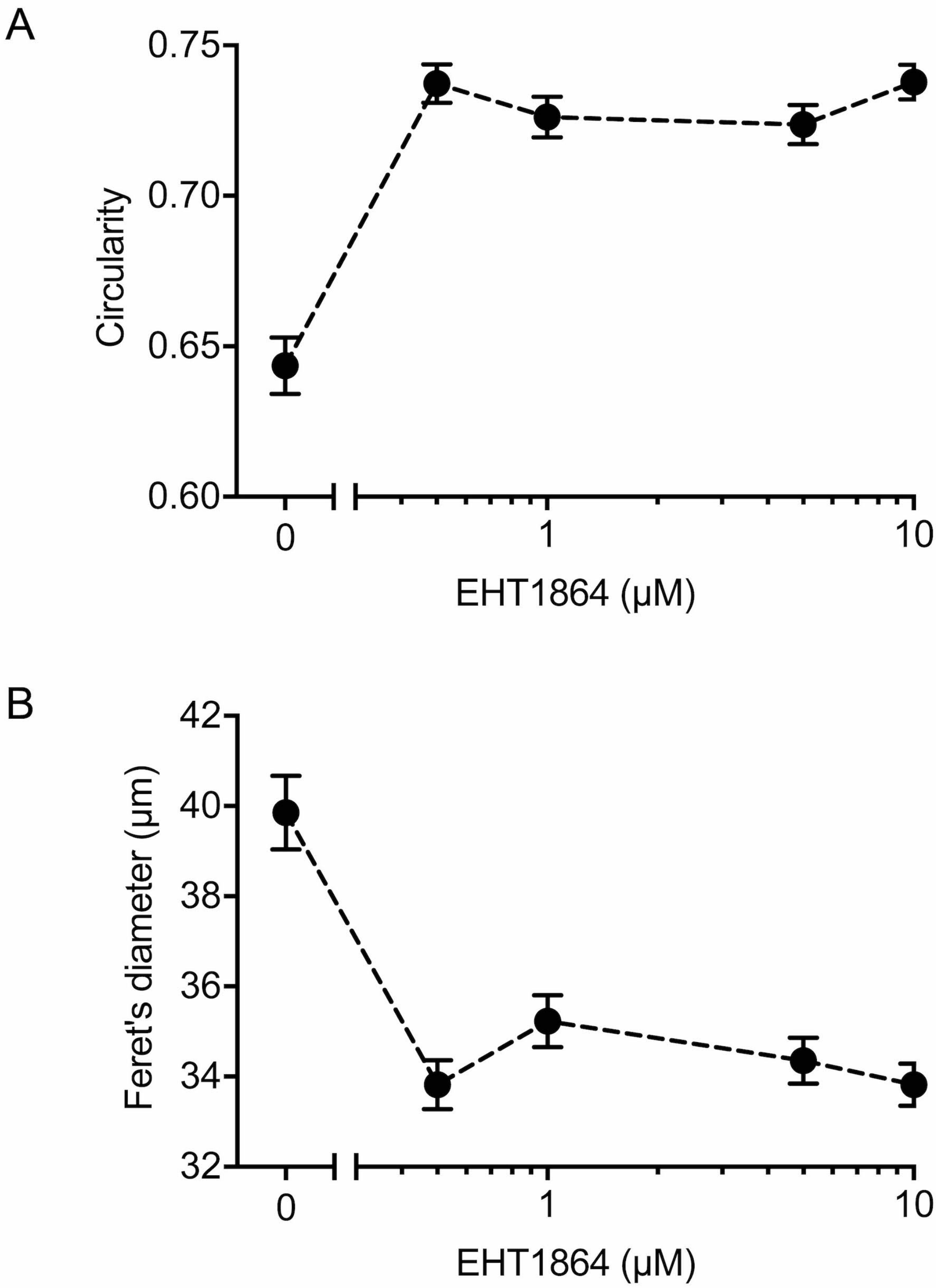
A
0.75
0.70
0.65
0.60
Circularity
0
1
10
EHT1864 (μM)
B
42
40
38
36
34
32
Feret's diameter (μm)
0
1
10
EHT1864 (μM)
